# The DNA replication initiation protein DnaD is recruited to a specific strand of the *Bacillus subtilis* chromosome origin

**DOI:** 10.1101/2022.04.18.488654

**Authors:** Charles Winterhalter, Simone Pelliciari, Daniel Stevens, Stepan Fenyk, Elie Marchand, Nora B Cronin, Panos Soultanas, Tiago R. D. Costa, Aravindan Ilangovan, Heath Murray

## Abstract

Genome replication is a fundamental biological activity shared by all organisms. Chromosomal replication proceeds bidirectionally from origins, requiring the loading of two helicases, one for each replisome. The molecular mechanisms for helicase loading at bacterial chromosome origins (*oriC*) are unclear. Here we investigated the essential DNA replication initiation protein DnaD in the model organism *Bacillus subtilis.* A set of DnaD residues required for ssDNA binding was identified, and photo-crosslinking revealed that this ssDNA binding region interacts preferentially with one strand of *oriC.* Biochemical and genetic data support the model that DnaD recognizes a new single-stranded DNA (ssDNA) motif located in *oriC* (*D*naD *R*ecognition *E*lement, “DRE”). Considered with cryo-electron microscopy (cryo-EM) imaging of full length DnaD, we propose that the location of the DRE within the *oriC* orchestrates strand-specific recruitment of helicase to achieve bidirectional DNA replication. These findings significantly advance our mechanistic understanding of bidirectional replication from a bacterial chromosome origin.

## INTRODUCTION

Faithful chromosome replication is universally essential to sustain life and initiates from specific loci called origins. Although the mechanisms triggering the onset of DNA replication are distinct across the phyla of life, all species require the loading of unwinding machines termed helicases prior to duplication of the genetic material (1). In bacteria, replication initiation generally occurs from a single origin (*oriC*) and proceeds bidirectionally towards an endpoint region called the terminus, typically located equidistance from the origin on the circular chromosome (2–5). To initiate this process, two toroid hexameric helicases need to be loaded at the origin, one around each strand (6). Despite fifty years of evidence for bidirectional DNA replication initiation in bacteria (7–9), the molecular mechanism for dual helicase loading at *oriC* remains unclear (10).

In *Bacillus subtilis*, DNA replication is initiated by five proteins that are sequentially recruited to the chromosome origin (Fig. 1A): DnaA, DnaD, DnaB and the DnaC:DnaI complex (helicase and AAA+ (*A*TPase *A*ssociated with various cellular *A*ctivities) loader, respectively) (11). The ubiquitous bacterial initiator DnaA binds at *oriC* and promotes open complex formation to allow helicase loading. DnaA is recruited to the origin by a conserved helix-turn-helix motif in domain IV, mediating binding to asymmetric double-strand DNA (dsDNA) sequences called DnaA-boxes (consensus 5′-TTATCCACA-3′) (12, 13). This promotes formation of a DnaA^ATP^ oligomer through extensive contacts between the protein’s AAA+ motifs, allowing it to engage a single strand of the DNA duplex (14) at specific trinucleotide motifs called DnaA-trios (consensus 3′-GAT-5′) (15–17).

**Figure 1.**
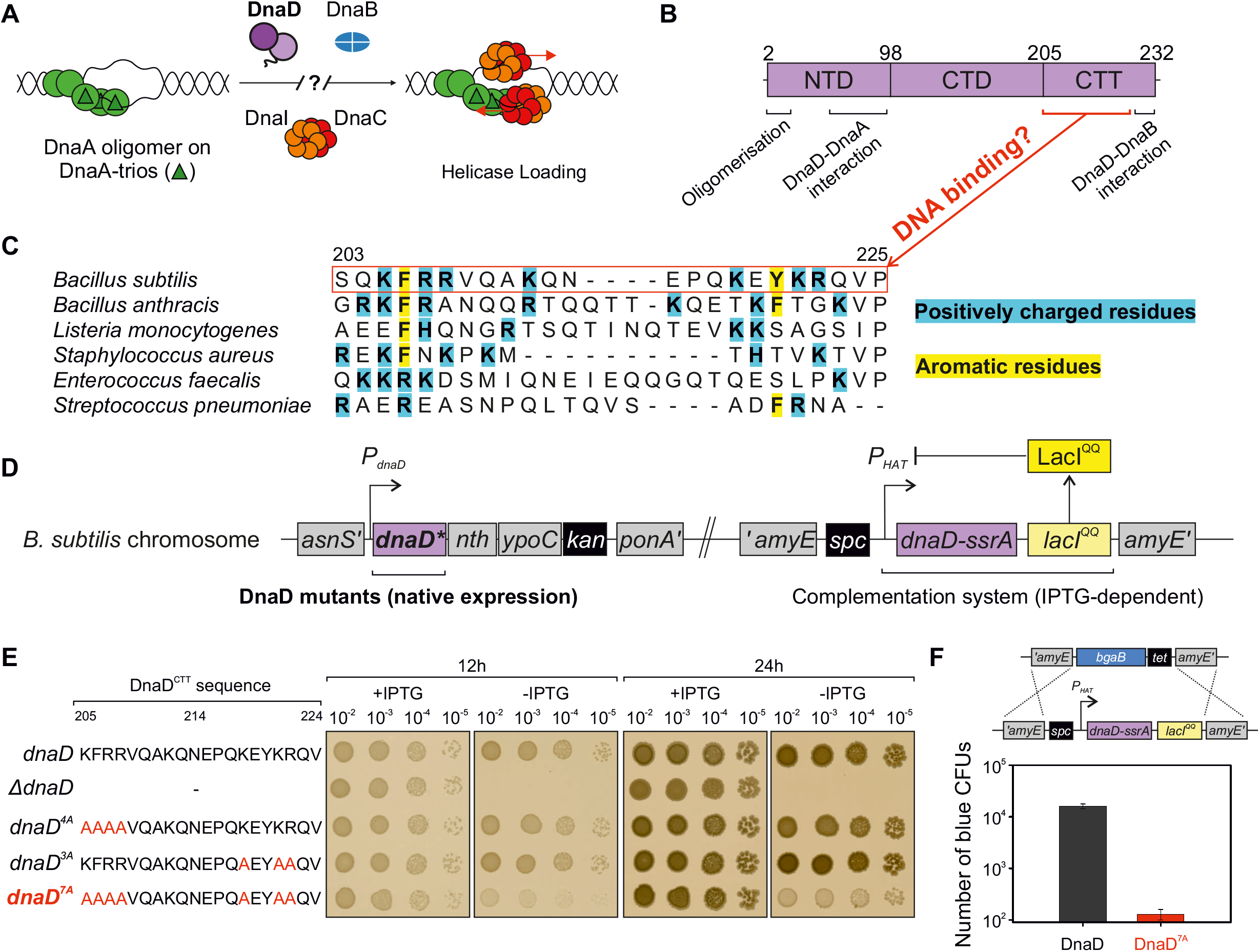
Identification of important DNA binding residues in *B. subtilis* DnaD. **(A)** Schematics of the helicase loading pathway in *B. subtilis* showing sequential recruitment of DnaA, DnaD, DnaB and the helicase complex DnaI-DnaC. **(B)** DnaD primary structure mapped with regions that are important for oligomerisation and interactions with DnaA and DnaB. NTD denote the N-Terminal Domain, CTD the C-Terminal Domain and CTT the C-Terminal Tail of DnaD. Amino acid sequence information shown in panel (C) suggested a potential DNA binding interface. **(C)** Protein sequence alignment of DnaD homologs showing the recurrence of positively charged (cyan) and aromatic residues (yellow) within the DnaD^CTT^. **(D)** Schematics of the complementation system used to screen the function of DnaD mutants in *B. subtilis*. Mutants are integrated at the endogenous *dnaD* locus and an inducible ectopic copy of *dnaD* (*dnaD-ssrA*) allows functional complementation by the addition of IPTG to the culture medium. The function of DnaD mutants can be determined upon removal of IPTG leading to repression of the *dnaD-ssrA* copy and degradation of DnaD-SsrA. **(E)** Spot titre analysis showing that multiple DnaD^CTT^ substitutions targeting residues highlighted in panel (C) were required to produce a growth phenotype as observed in *dnaD^7A^*. The presence or absence of IPTG indicates the induction state of the ectopic *dnaD-ssrA* cassette. *dnaD* (CW162), Δ*dnaD* (CW197), *dnaD^4A^* (CW412), *dnaD^3A^* (CW415), *dnaD^7A^* (CW647). (F) Quantification of the number of blue colony forming units (CFUs) obtained after attempting to remove the *dnaD-ssrA* cassette from a strain carrying wild-type DnaD or the 7A allele at the endogenous locus. Sequencing of eventual blue transformants in the *dnaD^7A^* background revealed that wild-type *dnaD* was restored. Recipient strains: DnaD (CW162), DnaD^7A^ (CW647).

A direct interaction between the AAA+ motifs of DnaA and of a helicase loader protein (*Aquifex aeolicus* DnaC) have been proposed to orchestrate loading of one helicase onto the strand bound by the DnaA oligomer (Fig. 1A) (18). However, the mechanism for loading a helicase onto the opposite strand is unclear. Recently, a region of DnaA domain I was identified to be essential for the recruitment of DnaD to *oriC* (19), raising the possibility that this interaction might guide the deposition of a second helicase at *oriC*.

DnaD comprises an N-terminal domain (DnaD^NTD^), a C-terminal domain (DnaD^CTD^) and a C-terminal tail (DnaD^CTT^). A recent alanine scan characterised the regions of DnaD necessary for its functions *in vivo*, including tetramerization and protein:protein interactions with DnaA and DnaB (Fig. 1B) (19). Conspicuously, this analysis did not identify any residues required for DNA binding. Previous studies indicated that DnaD displays a general affinity for DNA (20–25), with the ability to untwist dsDNA at high protein concentrations (23, 24). Furthermore, analysis of DnaD truncations *in vitro* strongly suggested that the DNA binding activity of DnaD was located in the C-terminal tail (20–22). Nonetheless, because specific mutations inactivating the DNA binding activity of DnaD have not been identified, the physiological relevance of DnaD binding to DNA is uncertain.

Here we identify DnaD variants specifically defective for binding DNA. Characterization of these DnaD proteins indicates that DnaD is recruited to a specific strand of *oriC* via a new ssDNA binding motif, the *D*naD *R*ecognition *E*lement (DRE). The location of the DRE opposite the DnaA-trios suggests a mechanism for directing strand-specific helicase loading to achieve bidirectional DNA replication initiation at the *B. subtilis* chromosome origin.

## MATERIALS AND METHODS

### Reagents

Nutrient agar (NA; Oxoid) was used for routine selection and maintenance of both *B. subtilis* and *E. coli* strains. Supplements were added as required: ampicillin (100 µg/ml), chloramphenicol (5 µg/ml), kanamycin (5 µg/ml), spectinomycin (50 µg/ml in *B. subtilis*, 100 µg/ml in *E. coli*), tetracycline (10 µg/ml), erythromycin (1 µg/ml) in conjunction with lincomycin (25 µg/ml), X-gal (0.01% v/v), IPTG (0.1 mM unless indicated otherwise). All chemicals and reagents were obtained from Sigma-Aldrich unless otherwise noted. Antibodies were purchased from Eurogentec. Plasmid extractions were performed using Qiagen miniprep kits. Other reagents used for specific techniques are listed within the method details.

### Biological resources: B. subtilis strains

*B. subtilis* strains are listed in Table S1 and were propagated at 37°C in Luria-Bertani (LB) medium unless stated otherwise in method details. Transformation of competent *B. subtilis* cells was performed using an optimized two-step starvation procedure as previously described (26, 27). Briefly, recipient strains were grown overnight at 37°C in transformation medium (Spizizen salts supplemented with 1 μg/ml Fe-NH_4_-citrate, 6 mM MgSO_4_, 0.5% w/v glucose, 0.02 mg/ml tryptophan and 0.02% w/v casein hydrolysate) supplemented with IPTG where required. Overnight cultures were diluted 1:17 into fresh transformation medium supplemented with IPTG where required and grown at 37°C for 3 hours with continual shaking. An equal volume of prewarmed starvation medium (Spizizen salts supplemented with 6 mM MgSO_4_ and 0.5% w/v glucose) was added and the culture was incubated at 37°C for 2 hours with continual shaking. DNA was added to 350 μl cells and the mixture was incubated at 37°C for 1 hour with continual shaking. 20-200 μl of each transformation was plated onto selective media supplemented with IPTG where required and incubated at 37°C for 24-48 hours. The genotype of all chromosomal origin and *dnaD* mutants was confirmed by DNA sequencing. Descriptions, where necessary, are provided below.

DnaD alanine substitution strains were generated by a blue/white screening assay using CW197 as parental strain and mutant plasmids obtained after Quickchange mutagenesis and sequencing as recombinant DNA (19). X-gal 0.016% w/v was added to transformation plates for detection of β-galactosidase activity and selection of kanamycin resistant white colonies that integrated mutant DNA by double-recombination. Three individual white colonies per mutant were then restreaked onto a medium either with or without IPTG to identify alleles of interest.

### Biological resources: E. coli strains and plasmids

*E. coli* transformations were performed in CW198 via heat-shock following the Hanahan method (28) for plasmids harbouring *dnaD* and propagated in LB with appropriate antibiotics at 37°C unless indicated otherwise in method details. Plasmids are listed in the Table S2 (sequences are available upon request). DH5α [F^-^ Φ80*lac*ZΔM15 Δ(*lac*ZYA-*arg*F) U169 *rec*A1 *end*A1 *hsd*R17(r_k_^-^, m_k_^+^) *pho*A *sup*E44 *thi*-1 *gyr*A96 *rel*A1 λ^-^] (29) was used for other plasmids construction. Descriptions, where necessary, are provided below.

pCW123, pCW310, pCW313 and pCW407 was generated by Quickchange mutagenesis using the oligonucleotides listed in Table S3.

pCW4 was generated by cloning *HindIII-SphI* PCR fragments generated using oligonucleotides listed in Table S3.

pCW66, pCW137, pCW171 were generated by ligase-free cloning via two-step assembly processes using oligonucleotides listed in Table S3. The underlined part of each primer indicates the region used to form an overlap. FastCloning (30) was used with minor modifications. PCR products (15 μl from a 50 μl reaction) were mixed and then subjected to a heating/cooling regime: two cycles at 98°C for 2 minutes then 25°C for 2 minutes, followed by one cycle at 98°C for 2 minutes and a final cycle at 25°C for 60 minutes. After cooling, *DpnI* restriction enzyme (1 μl) was added to digest parental plasmids and the mixtures were incubated at 37°C for ∼4 hours. Following digestion 10 μl of the PCR mixture was transformed into chemically competent *E. coli*. Where several primer pairs are listed for the construction of a single plasmid (Multi-step assembly column in Table S3), multiple rounds of ligase free cloning were performed to obtain the final constructs.

### Biological resources: oligonucleotides

All oligonucleotides were purchased from Eurogentec. Oligonucleotides used for plasmid construction are listed in Table S3 and oligonucleotides used for qPCR are listed in Table S4.

### Statistical analyses

Statistical analysis was performed using Student’s t-tests and p-values are given in figure legends. The exact value of *n* is given in method details and represents the number of biological repeats for an experiment. Tests were based on the mean of individual biological replicates and error bars indicate the standard error of the mean (SEM) across these measurements. Differences were considered as significant if their associated p-value was below 0.05. For ChIP experiments showing DnaA and DnaD depletion from the origin, the standard error of the mean was propagated by addition of the error from individual terms.

### Microscopy

To visualize cells by microscopy during the exponential growth phase, starter cultures were grown in imaging medium (Spizizen minimal medium supplemented with 0.001 mg/mL ferric ammonium citrate, 6 mM magnesium sulphate, 0.1 mM calcium chloride, 0.13 mM manganese sulphate, 0.1% w/v glutamate, 0.02 mg/mL tryptophan) with 0.5% v/v glycerol, 0.2% w/v casein hydrolysate and 0.1 mM IPTG at 37°C. Saturated cultures were diluted 1:100 into fresh imaging medium supplemented with 0.5% v/v glycerol and 0.1 mM IPTG and allowed to grow for three mass doublings. For DnaD mutants, early log cells were then spun down for 5 minutes at 9000 rpm, resuspended in the same medium lacking IPTG and further incubated for 90 minutes before imaging. For origin mutants, cells were grown in PAB at 20°C up to mid-exponential phase and 200 µL cells were further incubated with DAPI (1 µg/mL) for 15 min. Bacterial membranes were imaged by adding Nile red (1 µg/mL) directly to the agarose pad.

Cells were mounted on ∼1.4% agar pads (in sterile ultrapure water) and a 0.13- to 0.17-mm glass coverslip (VWR) was placed on top. Microscopy was performed on an inverted epifluorescence microscope (Nikon Ti) fitted with a Plan Apochromat Objective (Nikon DM 100x/1.40 Oil Ph3). Light was transmitted from a CoolLED pE-300 lamp through a liquid light guide (Sutter Instruments), and images were collected using a Prime CMOS camera (Photometrics). The fluorescence filter sets were from Chroma: GFP (49002, EX470/40 (EM), DM495lpxr (BS), EM525/50 (EM)), mCherry (49008, EX560/40 (EM), DM585lprx (BS), EM630/75 (EM)) and DAPI (49000, EX350/50, DM400lp, EM460/50). Digital images were acquired using METAMORPH software (version 7.7) and analysed using Fiji software (31). All experiments were independently performed at least twice, and representative data are shown.

The number of origins was quantified using the Trackmate plugin within the Fiji software (32). Background was subtracted from fluorescence images set to detect 8-10 pixel blob diameter foci over an intensity threshold of 150 relative fluorescence units. A mask containing the detected origin foci was created and merged with the nucleoids channel, and the number of origins per nucleoid was determined and averaged for a minimum of 100 cells from each strain that was examined. Because the nucleoid signal was more heterogenous in the presence of DnaD^7A^, we normalised individual nucleoid fluorescence by the average intracellular fluorescence observed in each microscopy image (Fig. S1). The count of nucleoids was determined using line plots of a 10 pixel width drawn across the length of cells. For each field of view, an average of the whole cell fluorescence was measured and used to normalise individual line plots, and the exact number of nucleoids per cell was assessed by counting the number of peaks crossing the zero line. This analysis was performed for at least two individual biological repeats.

### Phenotype analysis of dnaD mutants using the inducible dnaD-ssrA strain

Strains were grown for 18 hours at 37°C on NA plates unless otherwise stated (spot-titre assays) or in Penassay Broth (PAB, plate reader experiments) either with or without IPTG (0.1 mM unless otherwise stated). All experiments were independently performed at least twice and representative data are shown.

### Immunoblot analysis

Proteins were separated by electrophoresis using a NuPAGE 4-12% Bis-Tris gradient gel run in MES buffer (Life Technologies) and transferred to a Hybond-P PVDF membrane (GE Healthcare) using a semi-dry apparatus (Bio-rad Trans-Blot Turbo). DnaA, DnaD and FtsZ were probed with polyclonal primary antibodies (Eurogentec) and then detected with an anti-rabbit horseradish peroxidase-linked secondary antibody (A6154, Sigma) using an ImageQuant LAS 4000 mini digital imaging system (GE Healthcare). Detection of DnaA, DnaD and FtsZ was within a linear range. Experiments were independently performed at least twice and representative data are shown.

### Cryo-EM Sample Preparation and Data Collection

Four-microliter samples of purified wild-type DnaD were applied to plasma-cleaned Ultrafoil 2/2 200 grids, followed by plunge-freezing in liquid ethane using a Leica EM GP. Data collection was carried out at liquid nitrogen temperature on a Titan Krios microscope (Thermo Fisher Scientific) operated at an accelerating voltage of 300 kV. Micrograph movies were collected using EPU software (FEI) on a Gatan K3 detector in counting mode with a pixel size of 0.67 Å. A total of 3095 movie frames were acquired with a defocus range of approximately -0.7 to -2.7 μm. Each movie consisted of a movie stack of 30 frames with a total dose of 50 electron/Å^2^ over 1.5 seconds with a total dose of ∼50 electron/Å^2^ at a dose rate of 15 electron/pixel/second.

### Cryo-EM Image Processing, reconstruction and model fitting

The movie stacks were aligned and summed with dose-weighting using MotionCor2 (33). Contrast transfer function (CTF) was estimated by CtfFind4 (34), and images with poor CTF estimation were eliminated. A small subset of 200 micrographs were used to pick the particles using the blob picker tool in CryoSparc v3.1.0 and extracted with a box size of 400 pixels (35). These particles were 2D classified to generate a template which was subsequently used for particles picking using the template picket tool in CryoSparc. A total of 1299061 initial particles were picked using a box size of 400 x 400 pixels and subjected to several iterative rounds of 2D classification, removing particles belonging to poor template classes after each round of classification. A visual inspection of the 2D classes pointed to a clear two-fold symmetry within the particles. The final set of good particles (73190) selected after several iterative rounds of 2D classification were used to generate an *ab-initio* model using a C2 symmetry. Using the *ab-initio* model, the particles were subjected to 3D homogenous refinement using a dynamic mask between 6-14 Å resolutions where masking was at resolution higher than 12 Å in CryoSparc, which yielded a map at 10.1 Å with a conservative 0.5 FSC cut off.

A poly alanine model of the crystal structures of the DnaD^NTD^ (PDB 2V79) and the DnaD^CTD^ (PDB 2ZC2) were used to dock into the cryo-EM map of DnaD. The C-terminal domain can be readily identified and was docked into the density using Chimera (36). The entire N-terminal domain could not be readily docked in to the cryo-EM density hence a flexible fitting approach was adopted. The β-hairpin density could be easily identified within the map which was docked in the density first. The remainder of the structure was manually fitted in the density using Coot (37). A single round of real space refinement was performed to remove any clashes and idealize the model using Phenix (38).

### Marker frequency analysis

Strains were grown in PAB overnight at 37°C and diluted 1:100 the next morning in PAB. Cells were allowed to grow for 4 hours at 37°C or incubated until they reached an optical density of 0.4 for cold-sensitive assays performed at 20°C. Five hundred microliter samples were harvested and immediately mixed with sodium azide (1% w/v final) to arrest growth and genome replication. Cultures were collected by centrifugation, the supernatant was discarded and pellets were flash frozen in liquid nitrogen before gDNA extraction via the DNeasy blood and tissue kit (Qiagen).

qPCR was performed using the Luna qPCR mix (NEB) to measure the relative amount of origin DNA compared to the terminus. PCR reactions were run in a Rotor-Gene Q Instrument (Qiagen) using serial dilutions of the DNA and spore DNA was used as a control. Oligonucleotide primers were designed to amplify *incC* (qSF19/qSF20) and the terminus (qPCR57/qPCR58), were typically 20–25 bases in length and amplified a ∼100 bp PCR product (Table S4). Individual *ori:Ter* ratios were obtained in three steps: first, every Ct value was normalised to 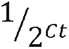, the dilution factor used during the qPCR and technical triplicates were averaged to a single enrichment value; second, origin enrichment was normalised by corresponding terminus values; third, *ori:Ter* values were normalised by the enrichment obtained for spore DNA. Error bars indicate the standard error of the mean for 2-4 biological replicates.

### Protein structure representations

Protein representations were generated using the Pymol Molecular Graphics 2.1 software (39), except for cryo-EM data and its associated model fitting that were performed in Chimera (36).

### Protein purification

Wild-type *dnaA* and *dnaD* were amplified by PCR using genomic DNA from *B. subtilis* 168CA and respectively cloned into pSF14 and pSF17 containing a His^14^-SUMO tag. DnaD mutants were created from pSF17 via quickchange reactions to introduce single or multiple substitutions. Wild-type DnaD expression was optimised by using a modified TIR sequence (pSP156) (40). Plasmids were propagated in *E. coli* DH5α and transformed in BL21(DE3)-pLysS for expression. Strains were grown in LB medium at 37°C. Overnight cultures were diluted 1:100 the next morning and at A_600_ of 0.6, 1 mM IPTG was added before further incubation at 30°C for 4 hours. Cells were harvested by centrifugation at 7000 g for 20 minutes, DnaA expression pellets resuspended in 40 ml of DnaA Ni^2+^ Binding Buffer (30 mM HEPES [pH 7.6], 250 mM potassium glutamate, 10 mM magnesium acetate, 30 mM imidazole), DnaD pellets in 40 ml of DnaD Ni^2+^ Binding Buffer (40 mM Tris-HCl [pH 8.0], 0.5 M NaCl, 5% v/v glycerol, 1 mM EDTA, 20 mM imidazole), each containing 1 EDTA-free protease inhibitor tablet (Roche #37378900) and then flash frozen in liquid nitrogen. Cell pellet suspensions were thawed and incubated with 0.5 mg/ml lysozyme on ice for 1h before disruption by sonication (1 hour at 20 W with 20 seconds pulses/rests intervals). Cell debris were removed from the lysate by centrifugation at 24,000 g for 30 minutes at 4°C, then passed through a 0.2 µm filter for further clarification. Further purification steps were performed at 4°C using a FPLC with a flow rate of 1 ml/min.

Clarified lysates were applied to a 1 ml HisTrap HP column (Cytiva). For DnaA, an additional wash with 10 ml Ni^2+^ High Salt Wash Buffer (30 mM HEPES [pH 7.6], 1 M potassium glutamate, 10 mM magnesium acetate, 30 mM imidazole) was performed. Materials bound to the column were washed with 10 ml of 10% Ni^2+^ Elution Buffer (DnaA: 30 mM HEPES [pH 7.6], 250 mM potassium glutamate, 10 mM magnesium acetate, 1 M imidazole; DnaD: 40 mM Tris-HCl [pH 8.0], 0.5 M NaCl, 1 mM EDTA, 0.5 M imidazole) and proteins were eluted with a 10 ml linear gradient (10-100%) of Ni^2+^ Elution Buffer. For DnaA, fractions containing the protein were applied to a 1 ml HiTrap Heparin HP affinity column (Cytiva) equilibrated in H Binding Buffer (30 mM HEPES [pH 7.6], 100 mM potassium glutamate, 10 mM magnesium acetate) and elution was carried out with a 20 ml linear gradient (20–100%) of H Elution Buffer (30 mM HEPES [pH 7.6], 1 M potassium glutamate, 10 mM magnesium acetate). Fractions containing proteins of interest were pooled and digested with 10 µl of 10 mg/ml His^14^-Tev-SUMO protease (41). For DnaD, digestion was performed at room temperature over the course of 48 hours and the same amount of His^14^-Tev-SUMO protease was added after 24 hours digestion.

Digestion reactions were applied to a 1 ml HisTrap HP column to capture non-cleaved monomers, His^14^-SUMO tag and His^14^-TEV-SUMO protease. Cleaved proteins were collected in the flow-through and their purity was confirmed using SDS-PAGE. Glycerol was added (DnaA: 20% v/v final; DnaD: 10% v/v final) and proteins aliquots were flash frozen in liquid nitrogen before being stored at -80°C.

### Bpa protein mutants expression and purification

Plasmids encompassing the non-canonical amino acid residue p-benzoyl-L-phenyl alanine (Bpa, Alfa Aesar #52083) were generated by inserting a *TAG* stop codon at the desired crosslinking position in the appropriate plasmid (construction detailed above). Electrocompetent *E. coli* BL21(DE3)-pLysS was co-transformed with 200ng of the Bpa-modified expression plasmid and pSup-BpaRS-6TRN(42), which encodes the necessary tRNA and tRNA synthase to recognize the *TAG* codon and to insert Bpa at this position. Co-transformed cells were grown at 37°C overnight on plates containing the appropriate antibiotics, and colonies were harvested the next day by directly scrapping them from the plate. Bpa was dissolved in 1M NaOH to obtain a 100 mM stock solution and then added to 2X YT media at 1 mM final concentration. After adjusting the pH of the solution to 7.0, cells collected from the transformation plate were inoculated at an optical density of 0.3, grown for 1h at 37°C and transferred to 30°C prior to addition of 1mM IPTG for protein expression. After 4h of vigorous shaking at 30°C, cultures were spun down, resuspended in the appropriate Ni^2+^ Binding Buffer (40 mM Tris-HCl [pH 8.0], 0.5 M NaCl, 5% v/v glycerol, 1 mM EDTA, 20 mM imidazole and one EDTA-free protease inhibitor tablet (Roche #37378900)), flash frozen in liquid nitrogen and purified as previously described.

### Bpa crosslinking

DnaD protein variants containing Bpa residues were adjusted to a final concentration of 0.1 µM and incubated with 30 nM of DNA probes in 1X strand separation buffer (10 mM HEPES-KOH (pH 7.6), 100 mM potassium glutamate, 5 mM magnesium acetate) for 10 minutes at room temperature. Reaction tubes were then transferred on ice and exposed to 365 nm wavelength light in a UV oven for 1 minute (∼400 mJ, Boekel #234100). A loading dye (4X SDS-page, Merck) was added to the samples to a final concentration of 1X and reactions were denaturated at 98°C for 5 minutes. Samples were loaded on a 4-12% Bis-Tris polyacrylamide gel, run in MES buffer for 45 min at 200 V and the gel was imaged using a Typhoon 9500 (GE healthcare, 600 V power).

### SEC-MALS

Experiments were conducted on a system comprising a Wyatt HELEOS-II multi-angle light scattering detector and a Wyatt rEX refractive index detector linked to a Shimadzu HPLC system (SPD-20A UV detector, LC20-AD isocratic pump system, DGU-20A3 degasser and SIL-20A autosampler) and the assays performed at 20°C. Solvent was 0.2 µm filtered before use and a further 0.1 µm filter was present in the flow path. The column was equilibrated with at least 2 column volumes of 40 mM Tris-HCl [pH 8], 500 mM NaCl, 1 mM EDTA, 20 mM imidazole, 2.5% v/v glycerol before use and flow was continued at the working flow rate until baselines for UV, light scattering and refractive index detectors were all stable.

The sample injection volume was of 100 µl, the Shimadzu LabSolutions software was used to control the HPLC and the Astra 7 software for the HELEOS-II and rEX detectors. The Astra data collection was 1 minute shorter than the LC solutions run to maintain synchronisation. Blank buffer injections were used as appropriate to check for carry-over between sample runs. Data were analysed using the Astra 7 software. Molecular weights were estimated using the Zimm fit method with degree 1 and a value of 0.179 was used for protein refractive index increment (dn/dc).

### Size-exclusion chromatography

SEC was performed with a Superdex 200 Increase 10/300GL column (Cytiva) at 4°C using a FPLC with a flow rate of 0.75 ml/min. The column was washed with two column volumes of MilliQ water before equilibration with 2.2 column volumes of DnaD buffer without glycerol (40 mM Tris-HCl [pH 8], 500 mM NaCl, 1 mM EDTA, 20 mM imidazole). Proteins were spun down at 17,000 g for 2 minutes to remove aggregates and 500 µl samples were applied to the column. A total of 1.5 column volumes of DnaD buffer without glycerol was passed through the column and data was extracted via the Unicorn 7 software.

### Sequence alignments

Multiple protein sequence alignments were performed using the Clustal Omega tool (43). DNA sequences were aligned via MUSCLE (44).

### ChIP-qPCR

Chromatin immunoprecipitation and quantitative PCR were performed as previously described (45) with minor modifications detailed below.

Strains were grown overnight at 30°C in Spizizen salts supplemented with tryptophan (20 µg/ml), glutamate (0.1% w/v), glucose (0.5% w/v) and casamino acid (0.2% w/v). The following day cultures were diluted 1:100 into fresh medium and allowed to grow to an A_600_ of 0.4. Samples were resuspended in PBS and cross-linked with formaldehyde (final concentration 1% v/v) for 10 minutes at room temperature, then quenched with 0.1 M glycine. Cells were pelleted at 4°C, washed three times with ice-cold PBS (pH 7.3) then frozen in liquid nitrogen and stored at -80°C. Frozen cell pellets were resuspended in 500 µl of lysis buffer (50 mM NaCl, 10 mM Tris-HCl pH 8.0, 20% w/v sucrose, 10 mM EDTA, 100 µg/ml RNase A, ¼ complete mini protease inhibitor tablet (Roche #37378900), 2000 K U/µl Ready-Lyse lysozyme (Epicentre)) and incubated at 37°C for 30 min to degrade the cell wall. 500 µl of immunoprecipitation buffer (300 mM NaCl, 100 mM Tris-HCl pH 7.0, 2% v/v Triton X-100, ¼ complete mini protease inhibitor tablet (Roche #37378900), 1 mM EDTA) was added to lyse the cells and the mixture was incubated at 37°C for a further 10 minutes before cooling on ice for 5 minutes. DNA samples were sonicated (40 amp) three times for 2 minutes at 4°C to obtain an average fragment size of ∼500 to 1000 base pairs. Cell debris were removed by centrifugation at 4°C and the supernatant transferred to a fresh Eppendorf tube. To determine the relative amount of DNA immunoprecipitated compared to the total amount of DNA, 100 µl of supernatant was removed, treated with Pronase (0.5 mg/ml) for 60 minutes at 37°C then stored on ice. To immunoprecipate protein-DNA complexes, 800 µl of the remaining supernatant was incubated with rabbit polyclonal anti-DnaA, anti-DnaD and anti-DnaB antibodies (Eurogentec) for 1 hour at room temperature. Protein-G Dynabeads (750 µg, Invitrogen) were equilibrated by washing with bead buffer (100 mM Na_3_PO_4_, 0.01% v/v Tween 20), resuspended in 50 µl of bead buffer, and then incubated with the sample supernatant for 1 hour at room temperature. The immunoprecipated complexes were collected by applying the mixture to a magnet and washed with the following buffers for 15 minutes in the respective order: once in 0.5X immunoprecipitation buffer; twice in 0.5X immunoprecipitation buffer + NaCl (500 mM); once in stringent wash buffer (250 mM LiCl, 10 mM Tris-HCl pH 8.0, 0.5% v/v Tergitol-type NP-40, 0.5% w/v sodium deoxycholate 10 mM EDTA). Finally, protein-DNA complexes were washed a further three times with TET buffer (10 mM Tris-HCl pH 8.0, 1 mM EDTA, 0.01% v/v Tween 20) and resuspended in 100 µl of TE buffer (10 mM Tris-HCl pH 8.0, 1 mM EDTA). Formaldehyde crosslinks of both the immunoprecipitate and total DNA were reversed by incubation at 65°C for 16 hours in the presence of 1000 U Proteinase K (excess). The reversed DNA was then removed from the magnetic beads, cleaned using QIAquick PCR Purification columns (Qiagen) and used for qPCR analysis.

qPCR was performed using the Luna qPCR mix (NEB) to measure the amount of genomic loci bound to DnaA, DnaD and DnaB. PCR reactions were run in a Rotor-Gene Q Instrument (Qiagen) using serial dilutions of the immunoprecipitate and total DNA control as template. Oligonucleotide primers were designed to amplify *oriC* (qSF19/qSF20) and the non-specific locus *yhaX* (oWKS145/oWKS146 (46)), and were typically 20–25 bases in length and amplified a ∼100 bp PCR product (Table S4). Error bars indicate the standard error of the mean for 3 biological replicates.

### DNA scaffolds

If DNA duplexes were used, a 20 µl reaction was assembled with 10 µM of each oligonucleotide in Oligo Annealing Buffer (30 mM HEPES–KOH [pH 8], 100 mM potassium acetate and 5 mM magnesium acetate). Samples were heated in a PCR machine to 95°C for 5 min and then cooled by 1°C/min to 20°C before being held at 4°C prior to dilution to 1 µM in Oligo Annealing Buffer.

### Fluorescence polarisation

Fluorescein-labelled oligonucleotides were diluted on ice to 2.5 nM in DNA strand displacement buffer (10 mM HEPES-KOH [pH 7.5]; 1 mM magnesium acetate; 100 mM potassium glutamate) and 37.5 µl diluted products were dispensed in individual wells of a flat-bottom black polystyrene 96-well plate (Costar #CLS3694). DnaD proteins were diluted to 2 µM in DNA strand displacement buffer and seven to nine 2-fold dilutions were performed in the same buffer. A negative control containing no protein was used as background. Probes were allowed to equilibrate at 20°C and 12.5 µl of protein dilutions were mixed with DNA (1.875 nM final DNA concentration in 50 µl volume). Reactions were allowed to incubate for 10 min at 20°C and polarisation readings were obtained using a plate reader (BMG Clariostar) set at the same temperature. For each experiment, technical replicates were averaged and the background corresponding to each probe was subtracted from experimental values, thus reporting the specific DnaD DNA binding activity on a single substrate. Error bars indicate the standard error of the mean over two to five biological replicates.

### DNA strand separation assay

DNA scaffolds that contained one oligonucleotide labelled with BHQ2, one with Cy5 and one unlabelled (12.5 nM final concentration) were diluted in 10 mM HEPES-KOH (pH 8), 100 mM potassium glutamate, 2 mM magnesium acetate, 30% v/v glycerol, 10% v/v DMSO and 1 mM nucleotide (ADP or ATP). All the reactions were prepared on ice to ensure the stability of the DNA probe, then allowed to equilibrate at 20°C. DnaA was added to a final concentration of 650 nM to allow displacement of all the probes. DnaD and BSA were used at the same final concentration. Reactions were performed using a flat-bottom black polystyrene 96-well plate (Costar #CLS3694) in triplicate and fluorescence was detected every minute over 60 min with a plate reader (BMG Clariostar). For all reactions a negative control without protein was used as background. At each timepoint the average background value was subtracted from the experimental value, thus reporting the specific DnaA activity on a single substrate. Error bars indicate the standard error of the mean over three biological replicates.

### Phenotype analysis of origin mutants

Strains were grown for up to 72 hours at 20°C or 37°C on NA plates. All experiments were independently performed at least twice and representative data are shown.

## RESULTS

### Two clusters of residues within the DnaD^CTT^ are important for protein activity in vivo

It is established that DnaD has an affinity for DNA and previous studies employing protein deletions reported that this activity involves the C-terminal tail (20–22). DnaD is conserved in Firmicute pathogens and an alignment of homologous DnaD^CTT^ sequences indicated the recurrence of positively charged and aromatic residues within this region (Fig. 1C). Combined with the observation that alanine scanning mutagenesis did not identify residues required for DNA binding (19), we hypothesized that the DnaD^CTT^ contains a robust DNA binding motif.

To investigate the role of these positively charged DnaD^CTT^ residues *in vivo*, we replaced the endogenous *dnaD* with alleles encoding for multiple alanine substitutions. Because DnaD is essential, to construct mutations at the native locus we employed an ectopic complementation system composed of an IPTG-inducible copy of *dnaD* carrying a C-terminal degradation tag (*dnaD-ssrA*) (Fig. 1D). Plasmids encoding *dnaD* alanine mutants were constructed and transformed into a complementation strain of *B. subtilis*. Substitutions replacing two clusters of residues (DnaD^7A^) were required to elicit an observable growth defect as revealed by spot titre analysis (Fig. 1E). While the DnaD^7A^ variant was mildly underexpressed compared to the wild-type protein (Fig. S3A), we note that a Δ*dnaD* mutant strain can sustain growth even when the level of complementing DnaD-ssrA was undetectable by immunoblot (Fig. S2). Thus, it seems unlikely that changes in expression level can explain the phenotype of *dnaD^7A^*.

The titration of DnaD, showing that low protein levels could support cell growth, alerted us to the possibility that leaky expression of DnaD-ssrA may contribute to the viability of *dnaD^7A^*. To examine the activity of DnaD^7A^ in the absence of DnaD-ssrA, we attempted to replace the ectopic *dnaD-ssrA* with a cassette containing *tet* (tetracycline resistance) and *bgaB* (β-galactosidase) genes, using recipient strains expressing either wild-type DnaD or DnaD^7A^ from the endogenous locus. Selection for Tet^R^ showed a ∼100-fold reduction in the number of colonies obtained in the presence of *dnaD^7A^* (Figs. 1F and S3B). Screening for β-galactosidase activity revealed that only a minority of the transformants observed in the presence of *dnaD^7A^* correctly integrated the *bgaB* cassette (Fig. S3B), and DNA sequencing showed that these rare blue transformants had, in fact, restored wild-type *dnaD* at the native locus. To ensure that the *dnaD^7A^*strain was genetically competent, as a control we independently performed transformations using a DNA fragment containing an antibiotic cassette that integrates at an unlinked locus. Here a similar number of colonies was observed for each strain, thus the low frequency of transformants observed when attempting to replace *dnaD-ssrA* appears specific (Fig. S3C). Taken together, these genetic experiments indicate that two clusters of positively charged residues in the DnaD^CTT^ are essential for protein activity.

### The DnaD^CTT^ mutant DnaD^7A^ is defective in DNA replication initiation

In *B. subtilis*, DnaD has a role in both the replication initiation and restart pathways (46–48) and was also suggested to be involved in chromosome organisation via DNA remodelling activities (25, 49). To investigate the defects associated with the *dnaD^7A^* allele, we used fluorescence microscopy to visualise the impact of this variant on chromosome content. Here a fluorescence repressor/operator system was used to report the location of the chromosome origin (TetR-mCherry binding to a *tetO* array (50)) and the nucleoid was labelled with the nucleoid associated protein Hbs-GFP (Fig. 2A) (51). Images were taken following depletion of the ectopically expressed DnaD-ssrA. Under the growth conditions used, cells containing wild-type DnaD typically displayed a pair of chromosome origins per nucleoid (*ori:nuc*) (Fig. 2B-D) (52, 53). In contrast, cells expressing DnaD^7A^ contained a lower number of *oriC* per nucleoid (Fig. 2D). This suggests that mutations in the DnaD^CTT^ impact the onset of DNA replication.

**Figure 2.**
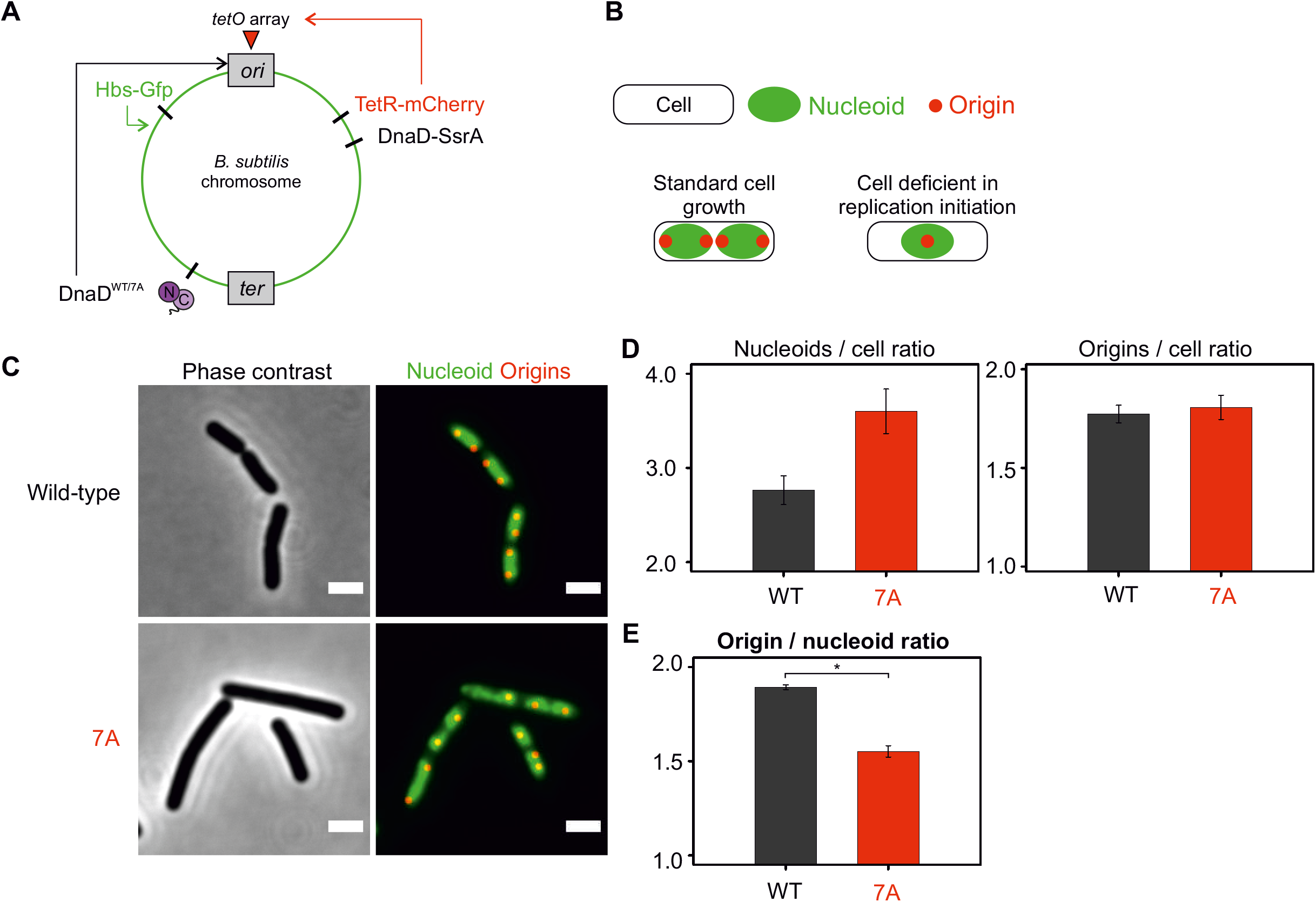
DnaD^7A^ has an aberrant growth phenotype and is depleted from the replication origin. **(A)** Schematics of the number of origins and nucleoids per cell showing the typical patterns associated with cells displaying a normal growth phenotype or cells that are deficient in replication initiation. **(B)** Schematics of the dual fluorescence system engineered to label chromosome origins and the nucleoid. TetR-mCherry (red fluorescence) is constitutively expressed and binds an array of *tetO* sites located near the origin of replication. Hbs-GFP (green fluorescence) is constitutively expressed, non-specifically binds double-stranded DNA and allows visualisation of the nucleoid. **(C)** Representative images of *dnaD* strains observed by fluorescence microscopy via the system described in panels (A-B). Red dots show chromosome origins and the green signal allows localisation of the nucleoid. Wild-type corresponds to a strain encoding the ectopic dnaD-ssrA cassette that was depleted from IPTG and relied on the endogenous copy of wild-type *dnaD* (WT) to sustain growth (CW517). 7A (CW667). Scale bar indicates 2μm. **(D-E)** Single-cell image analysis performed on the data obtained from experiments shown in panel (C). **(D)** shows the number of nucleoids per cell (left) and origins per cell (right). **(E)** shows the number of origins per nucleoid. Error bars show the standard error of the mean for at least two biological replicates where over 100 cells were studied for each cell background. * shows a p-value of 0.0091.

### DnaD binds ssDNA via the DnaD^CTT^

To directly investigate the DNA binding activity of DnaD *in vitro*, we established a fluorescence polarization assay to detect the interaction of DnaD with fluorescein labelled DNA substrates (Fig. 3A) (54). It was found that wild-type DnaD (i) binds ssDNA with a higher affinity than dsDNA (Fig. 3B; here the dsDNA probe corresponds to the annealing of ssDNA sequences), (ii) displays a preference for polythymidine (Fig. 3C), and (iii) requires a ssDNA substrate between 11-15 nucleotides in size (Fig. 3D).

**Figure 3.**
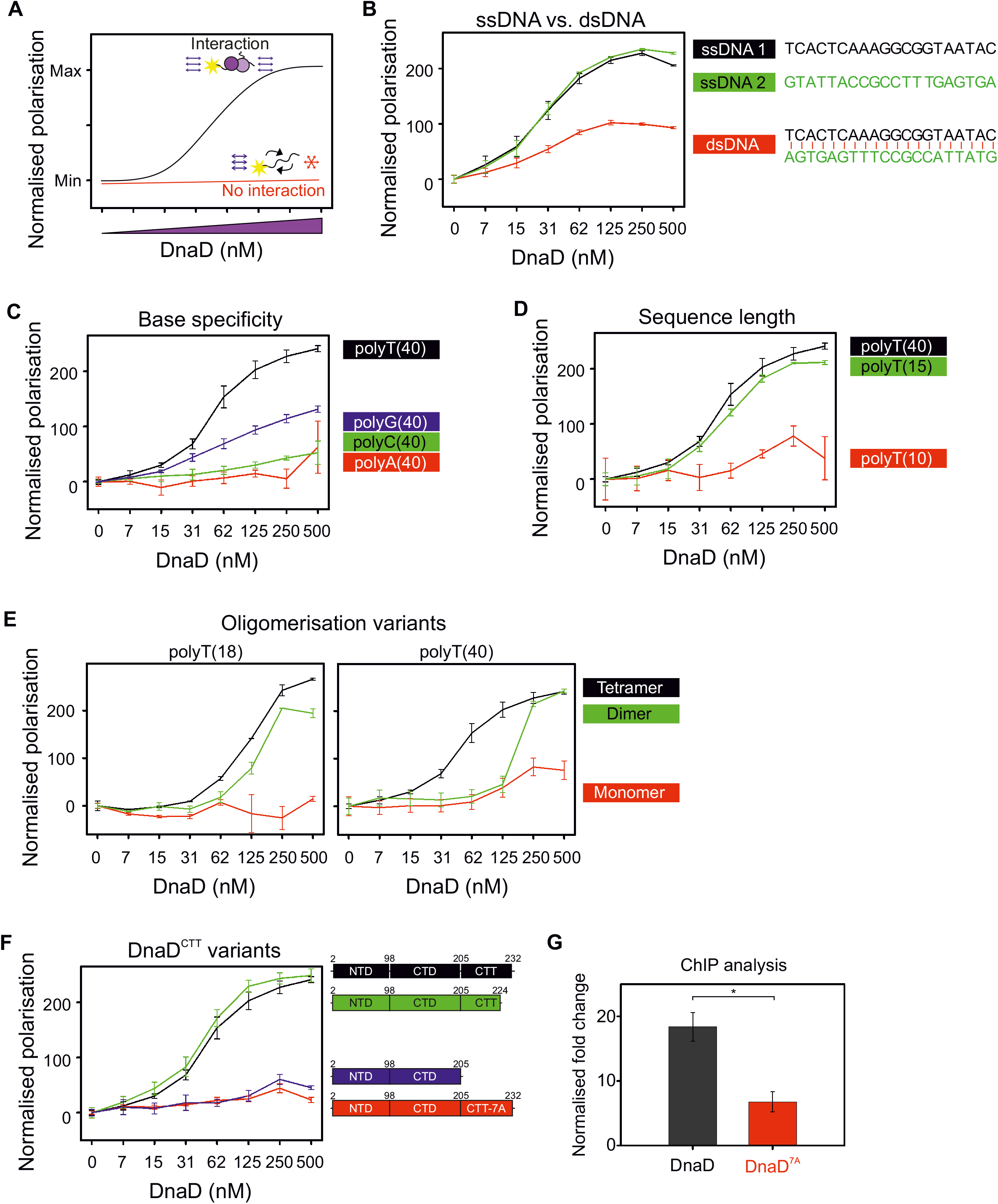
DnaD binds ssDNA through the DnaD^CTT^. **(A)** Schematics of a fluorescence polarisation assay showing interaction (black line) or no interaction (red line) between DnaD and DNA. In this assay, polarised light is shone upon a protein:DNA mixture. For a fixed concentration of fluorescein-labelled DNA, titration of DnaD produces a sigmoid curve where light shone upon samples remains polarised at high protein concentrations, suggesting an interaction between protein and DNA. If DNA is unbound and tumbles rapidly, polarised light shone upon the sample remains polarised and indicates that there is no protein:DNA interaction. A linear increase of polarisation values over titration of the protein indicates a non-specific interaction between the protein and fluorescein. **(B-E)** show fluorescence polarisation analyses of wild-type DnaD incubated with a range of DNA substrates. **(B)** shows that DnaD preferentially binds ssDNA over dsDNA. Sequences were randomly generated and the DNA duplex (dsDNA) corresponds to the annealing of individual ssDNA sequences. ssDNA 1 (oCW1036), ssDNA 2 (oCW1124) and dsDNA (oCW1034:oCW1124). **(C)** shows that DnaD preferentially binds thymidine bases. The protein was incubated with homopolymer sequences of a size of 40 nucleotides. polyT(40) (oCW837), polyG(40) (oCW854), polyC(40) (oCW852), polyA(40) (oCW853). **(D)** shows that DnaD preferentially binds substrates of a size equal to or over 15 nucleotides. Substrates were chosen based on the binding preference observed in panel (C). polyT(40) (oCW837), polyT(15) (oCW1029), polyT(10) (oCW834). **(E)** shows that the monomer mutant DnaD^L22A^ (red line) could not bind ssDNA and that the dimer variant DnaD^F6A^ (green line) and wild-type (black line) were both able to bind ssDNA. Polarisation profiles are shown for binding to a homopolymer of 18 thymidine nucleotides (oCW1088) or 40 thymidine nucleotides (oCW837). **(F)** Fluorescence polarisation analysis of DnaD protein variants indicates that DnaD residues 205-224 are important of DNA binding activity and that the DnaD^CTT^ variant DnaD^7A^ is defective in ssDNA binding. All proteins were incubated with a homopolymer of 40 thymidine bases (oCW837). The black line corresponds to wild-type DnaD, the green line to a variant lacking the last eight amino acid residues of the DnaD^CTT^, the blue line to a variant lacking the entire DnaD^CTT^ and the red line to the DnaD^7A^ mutant where multiple alanine substitutions were targeted at positively charged and aromatic residues of the DnaD^CTT^. Error bars in panels (B-E-F) indicate the standard error of the mean for 2-4 biological replicates. **(G)** ChIP analysis showing that the DnaD variant 7A is less enriched than the wild-type protein at *oriC*. The normalised fold change corresponds to specific enrichment where background of the DnaD-SsrA signal after IPTG depletion was subtracted from absolute values (signal from CW197). Primers used for the origin anneal within the *incC* region. * shows a p-value of 0.0129 (3 biological replicates). DnaD (CW162), DnaD^7A^ (CW647).

It has been observed that oligomerisation is crucial for DnaD function *in vivo*, and amino acid substitutions have been identified that specifically disrupt either dimerization (DnaD^L22A^) or tetramerization (DnaD^F6A^) (19). Using a dT_18_ substrate, it was observed that monomeric DnaD^L22A^ cannot bind, while dimeric DnaD^F6A^ and wild-type DnaD bound similarly (Fig. 3E left). Interestingly, the wild-type protein showed better binding affinity than DnaD^F6A^ to a dT_40_ substrate (Fig. 3E right, note that DnaD^L22A^ remained unable to bind this ssDNA). These results suggest that DnaD may have different modes of ssDNA binding and extends the findings from previous studies where DnaD was shown to bind polythymidine substrates (21) or other ssDNA substrates of variable size and sequence (20,23,55). However, because the DnaD tetramer was most active binding DNA *in vitro* and because tetramer formation appears to be essential *in vivo* (19), we suspect this binding activity is most critical.

Next, DnaD variants with alterations to the C-terminal tail were purified: DnaD^7A^ and two truncations, which removed either the DnaB interaction region (DnaD^1-224^ (19)) or the entire C-terminal tail containing the putative ssDNA binding residues (DnaD^1-205^). Both DnaD^7A^ and DnaD^1-205^ were unable to interact with a fluorescently labelled dT_40_ substrate, whereas DnaD^1-224^ retained activity similar to wild-type (Fig. 3F). Size exclusion chromatography (SEC) followed by multiple-angle light scattering (MALS) showed that DnaD^1-205^, carrying the largest truncation, was a tetramer (Fig. S4A-C). We also verified using SEC that DnaD^7A^ and DnaD^1-224^ assembled into tetramers (Fig. S4D). Taken together with the genetic analysis above, the results suggest that the essential activity of DnaD located between residues 205-224 is to bind ssDNA.

### The DnaD^CTT^ is required for DnaD recruitment to oriC

The replication initiation pathway in *B. subtilis* starts with binding of DnaA to the chromosome origin, followed by recruitment of DnaD (11). To determine whether the DnaD^7A^ protein was defective for DNA binding *in vivo*, we determined the amount of DnaD^7A^ recruited *to oriC* using chromatin immunoprecipitation followed by quantitative PCR (ChIP-qPCR). In the context of the *dnaD-ssrA* complementation system, we depleted DnaD-ssrA from exponentially growing cells for 90 minutes, crosslinked proteins to DNA and immunoprecipitated DnaD complexes. DnaD was enriched at *oriC* in cells that harboured either wild-type *dnaD*, *dnaD^7A^*, or Δ*dnaD* at the endogenous locus (Fig. S5A-C), indicating that DnaD-ssrA remained a detectable even following depletion (Fig. S5C-D). Nonetheless, assuming the level of DnaD-ssrA detected in the Δ*dnaD* strain represents the background, quantification of ChIP-qPCR data indicate that DnaD^7A^ is less enriched than wild-type at *oriC* (Figs. 3H). Combined with the *in vitro* analyses, the results suggest that the DnaD^CTT^ mediates DNA binding *in vivo*.

### Architecture of a DnaD dimer determined by cryo-electron microscopy

The structure of the DnaD^CTT^ has not been observed experimentally. In an attempt to place the DnaD^CTT^ within the DnaD tetramer, we characterized the full-length protein using single particle cryo-electron microscopy (cryo-EM, Table S5). Data analysis from 2D classes (Fig. 4A) and image processing revealed a 10 Å resolution map of a DnaD dimer (Figs. 4B and S6A). Although DnaD is a tetramer in solution (19), it was clear from the cryo-EM data that only a dimer of DnaD protomers could fit the observed density (Fig. 4B). We speculate that buffer conditions or cryo-EM sample preparation may destabilize DnaD tetramers, resulting in dissociation to stable dimers.

**Figure 4.**
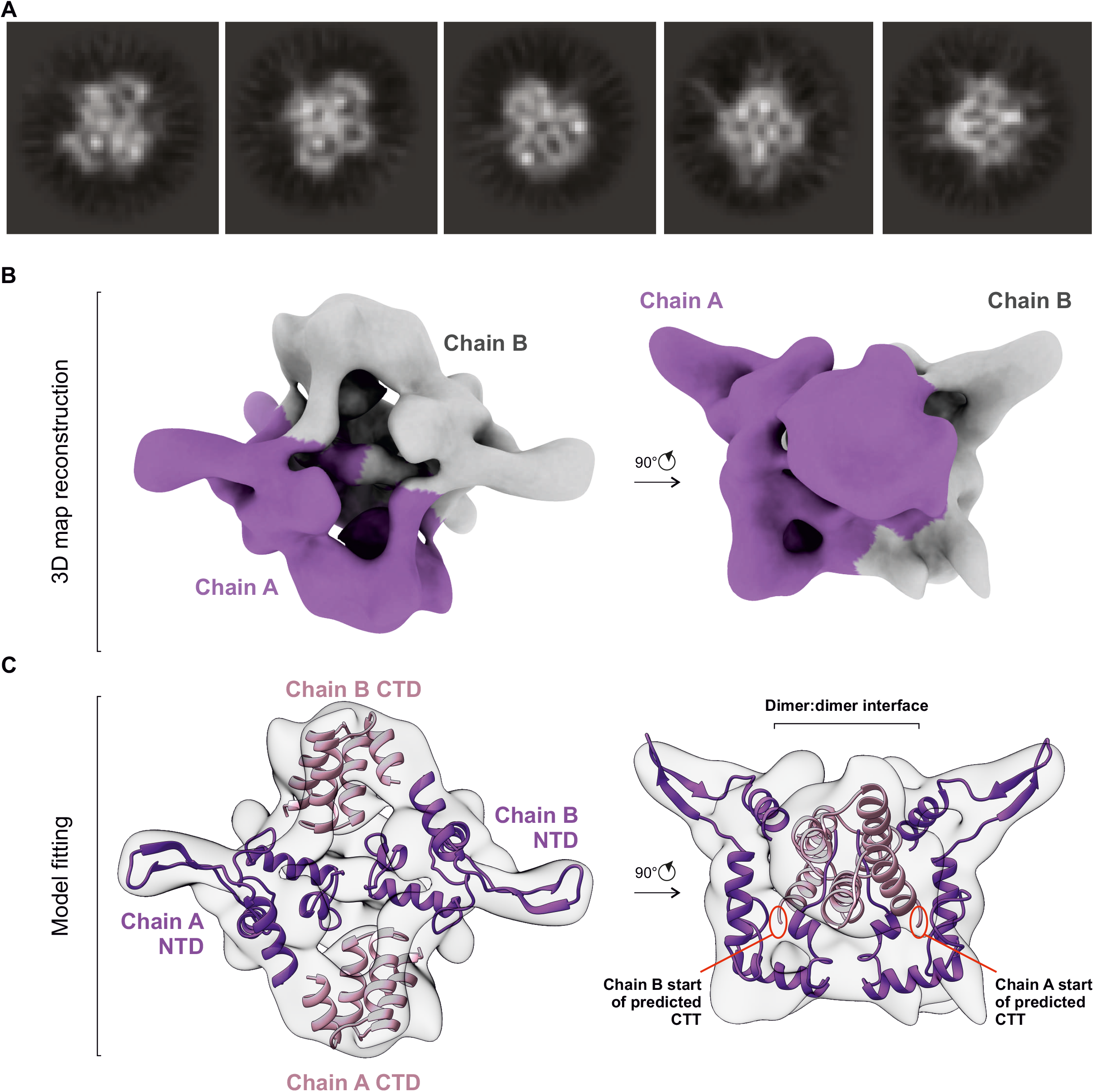
Cryo-EM identification of a DnaD dimer. **(A)** 2D classes observed by cryo-EM. **(B)** Reconstructed three-dimentional volume (density map) corresponding to a DnaD dimer. Each protomer is indicated by chain A and B labels shown in purple and grey, respectively. **(C)** Cryo-EM density map shown in panel (B) fitted with the available DnaD structures (purple N-terminal domains from PDB 2V79 and pink C-terminal domains from 2ZC2).

DnaD^CTD^ substructures and the β-hairpin within the DnaD^NTD^ were immediately identified within the cryo-EM map, and a poly alanine model of full-length DnaD encompassing a pair of DnaD^NTD^ and DnaD^CTD^ could be generated (Fig. 4C). While the previously published DnaD^CTD^ structure (PDB 2ZC2) agrees well with the cryo-EM model, some differences were observed with the arrangement of α-helices and β-strands described in the crystal structure of the DnaD^NTD^ (PDB 2v79) (56). Despite these alterations, it was noted that essential residues required for interacting with DnaA were found to run along one surface of the DnaD N-terminal domain, supporting the relevance of the cryo-EM structure (Fig. S6B).

Fitting NTD and CTD domains into the map suggests that the DnaD^CTD^ is positioned adjacent to the DnaD^NTD^. Critically, placement of the DnaD^CTD^ suggests that the flexible DnaD^CTT^ (which was not assigned within the map, consistent with previous studies (20)) would extend from a location distal to the proposed dimer:dimer interface (Fig. 4C), presumably making it available to bind ssDNA.

### DnaD interacts with a specific single-strand DNA binding element within the unwinding region of oriC

It has been found that DnaD requires a direct interaction with DnaA to be recruited to *oriC* (19). We speculated that this protein:protein interaction could direct DnaD to the *B. subtilis* chromosome origin for it to bind DNA (Fig. 5A) (57). To test for a direct physical interaction between the DnaD^CTT^ and *oriC*, we used protein:DNA photo-crosslinking. Here the non-natural photoactivatable amino acid p-benzoyl-L-phenylalanine (Bpa) was incorporated at the 221^st^ amino acid (DnaD^K221Bpa^, Fig. 5B-C) using an engineered amber suppressor tRNA with its cognate aminoacyl tRNA synthase (42). To detect DnaD:DNA crosslinking following UV irradiation, oligonucleotide substrates were fluorescently labelled and nucleoprotein complexes were separated by size using SDS-PAGE.

**Figure 5.**
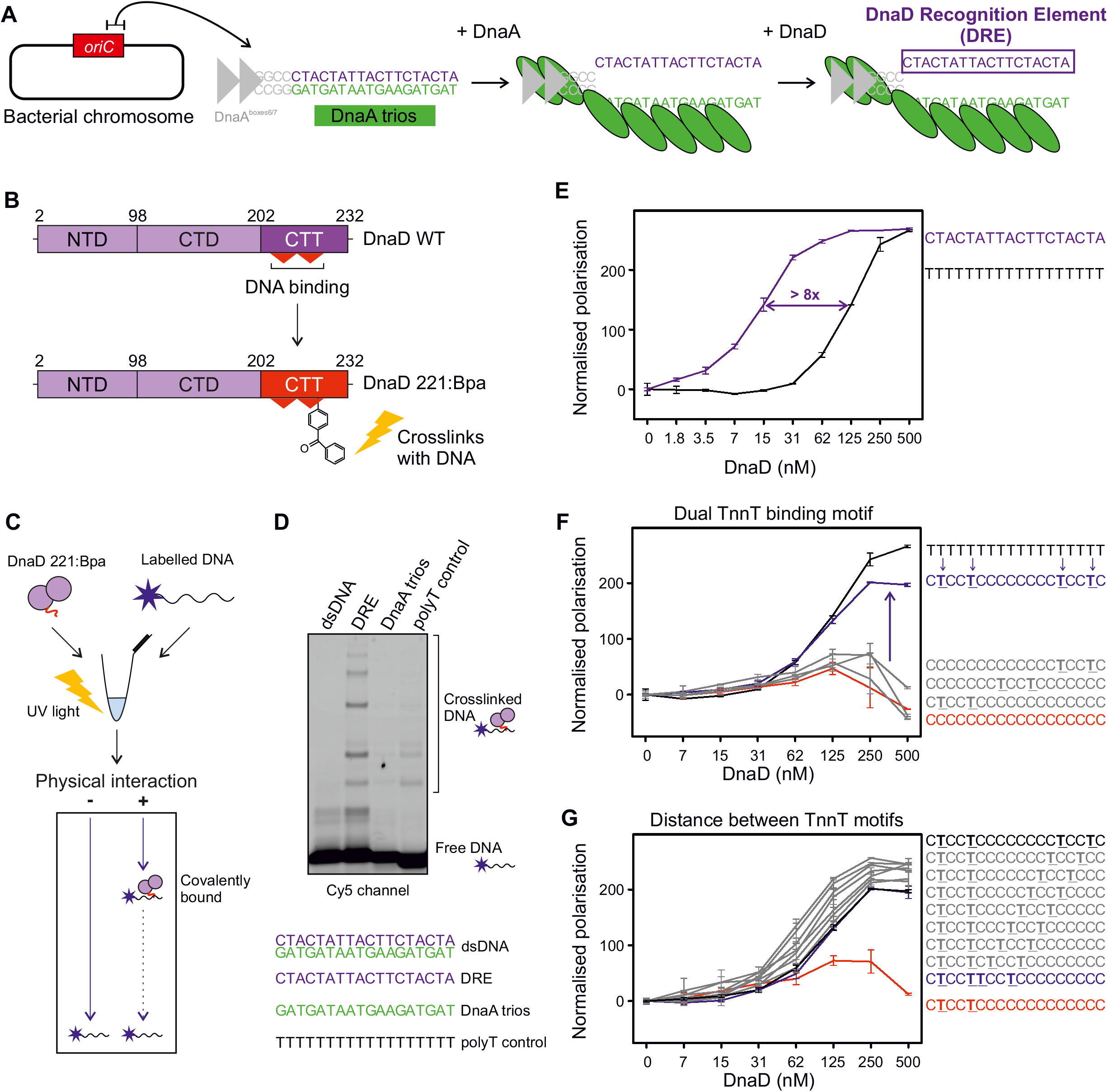
DnaD binds to the sequence facing the DnaA trios. **(A)** Illustration of the proposed basal origin unwinding mechanism in *B. subtilis*. The closed origin region *incC* is bound by DnaA on DnaA boxes, which leads to partial melting of the origin and strand separation via DnaA oligomer formation on the DnaA-trios. The complementary sequence to the DnaA-trios is proposed to be a specific binding substrate for DnaD. **(B)** DnaD primary structure schematics showing the incorporation of a non-natural amino acid residue within DnaD^CTT^ to probe physical interactions between DnaD and DNA. As opposed to the wild-type protein (WT), DnaD 221:Bpa can be crosslinked to DNA if the residue DnaD^221^ comes in contact with nucleic acids. NTD denote the N-Terminal Domain, CTD the C-Terminal Domain and CTT the C-Terminal Tail of DnaD. **(C)** Schematics of the Bpa crosslinking assay used to probe direct interactions between DnaD and DNA. DnaD 221:Bpa is incubated with a labelled oligonucleotide and reactions are crosslinked using UV exposure before running products on a denaturing gel. Following electrophoresis, free DNA runs to the bottom of the gel and the migration of covalently bound DNA:protein complexes is delayed. **(D)** Bpa crosslinking assay showing that DnaD specifically interacts with the DRE via the amino acid residue 221 in the DnaD^CTT^. Incubation with Cy-5 labelled oligonucleotides shows that DnaD interacts with ssDNA and has greater binding affinity to the DRE compared to the DnaD trios. Oligonucleotide sequences are indicated below the gel: dsDNA (oSP1132:oCW1040), DRE (oSP1132), DnaA trios (oSP1133), polyT control (oSP1135). **(E-G)** show fluorescence polarisation analyses of wild-type DnaD incubated with a range of DNA substrates. **(E)** shows that DnaD has significantly higher affinity for the DRE sequence than a polyT substrate of the same size. The purple line shows binding to the DRE (oCW1039) and black line binding to a polyT substrate (oCW1088). **(F)** shows that DnaD requires at least two TnnT elements to bind ssDNA. The black line shows binding to a polyT18 substrate (oCW1088), the red line shows incubation with a polyC18 substrate (oCW1089), the blue line indicates binding to a polyC substrate with two TnnT elements (oCW1128), and grey lines (from top to bottom) show incubation profiles with three oligonucleotides containing a single TnnT element (oCW1165, oCW1140 and oCW1164, respectively). **(G)** shows that the distance separating two TnnT elements does not affect the ability of DnaD to bind ssDNA. The black line shows binding to a polyC substrate with two TnnT elements separated by a distance of 8 nucleotides (oCW1128). Grey lines show binding of DnaD to various ssDNA sequences where the distance separating TnnT repeats in the context of a polyC substrate is gradually reduced (from top to bottom oCW1491, oCW1492, oCW1493, oCW1494, oCW1495, oCW1496 and oCW1497) down to juxtaposition of the two TnnT repeats (blue line, oCW1498). The red line shows the incubation profile of DnaD with a substrate containing only one TnnT element (oCW1164). Error bars in panels (E-G) indicate the standard error of the mean for 2-6 biological replicates.

A clear set of higher molecular weight species was detected when the stand complementing the DnaA-trios was incubated with DnaD^K221Bpa^ (Fig 5D). Annealing this strand with the DnaA-trios to form a dsDNA substrate abolished UV crosslinking (Fig. 5D), consistent with the model that DnaD prefers to bind ssDNA (Fig. 3B). Moreover, DnaD^K221Bpa^ crosslinking to this strand appeared most efficient, compared to ssDNA substrates with either DnaA-trios or a homopolymeric sequence (Figs. 5D, S7). Treatment with proteinase K degraded the higher molecular weight complexes, indicating that these are bonafide nucleoprotein complexes (Fig. S7). Finally, fluorescence polarization confirmed that DnaD binds with highest affinity to the strand complementing the DnaA-trios (Fig. 5E). Based on these observations and combined with the finding that DnaD binds the dT_18_ substrate better than other homopolymeric ssDNA (Figs. S7B and S8A), we hypothesized that thymidine might be a specificity determinant within the preferentially bound strand.

To identify putative DnaD binding motifs within ssDNA, thymidine bases were systematically introduced within inert dC_18_ or dA_18_ substrates and DnaD binding was assessed using fluorescence polarization. The results indicate that two motifs of 5’-TnnT-3’ are necessary and sufficient for DnaD to associate specifically with ssDNA (Figs. 5F and S8B-D). Intriguingly, the distance between the two 5’-TnnT-3’ motifs can be varied widely (Fig. 5G). This is consistent with both the high affinity of DnaD for the ssDNA complementary to the DnaA-trios (Fig. 5D-E) and the multiple higher molecular weight species detected by UV crosslinking, as it contains five potential 5’-TnnT-3’ motifs. It may be that binding of one DnaD protomer to a first 5’-TnnT-3’ motif enhances binding of another protomer to a second 5’-TnnT-3’ motif. Based on these properties, we have termed the ssDNA sequence complementary to the DnaA-trios the *D*naD *R*ecognition *E*lement (DRE), and we propose that pairs of 5’-TnnT-3’ motifs are critical for DnaD binding.

### The DRE is required for recruiting DnaD to the chromosome origin

We wanted to determine the role of the DRE *in vivo.* However, the DRE and the DnaA-trios are inherently linked ssDNA binding motifs that we propose are bound by DnaD and DnaA, respectively. Therefore, we set out to identify separation of function mutations within this region that support DnaA activities while disrupting DnaD binding.

Previous studies have suggested that the DnaA-trios closest to the DnaA-boxes are the most critical for DnaA unwinding activity (15-17,58). Therefore, we hypothesized that mutating the 5’-TnnT-3’ motif furthest from the DnaA-boxes might preferentially inhibit DnaD binding while leaving DnaA activity relatively unperturbed. Consistent with this notion, it was found that DnaA strand separation activity *in vitro* was similar to wild-type when the distal 5’-TnnT-3’ motif was changed to 5’-AnnA-3’ (Fig. 6A-B). Relatedly, we found that DnaD alone does not promote DNA strand separation (Figs. S9A-C) and that DnaD does not specifically stimulate DnaA strand separation activity (Figs. S9D-E).

**Figure 6.**
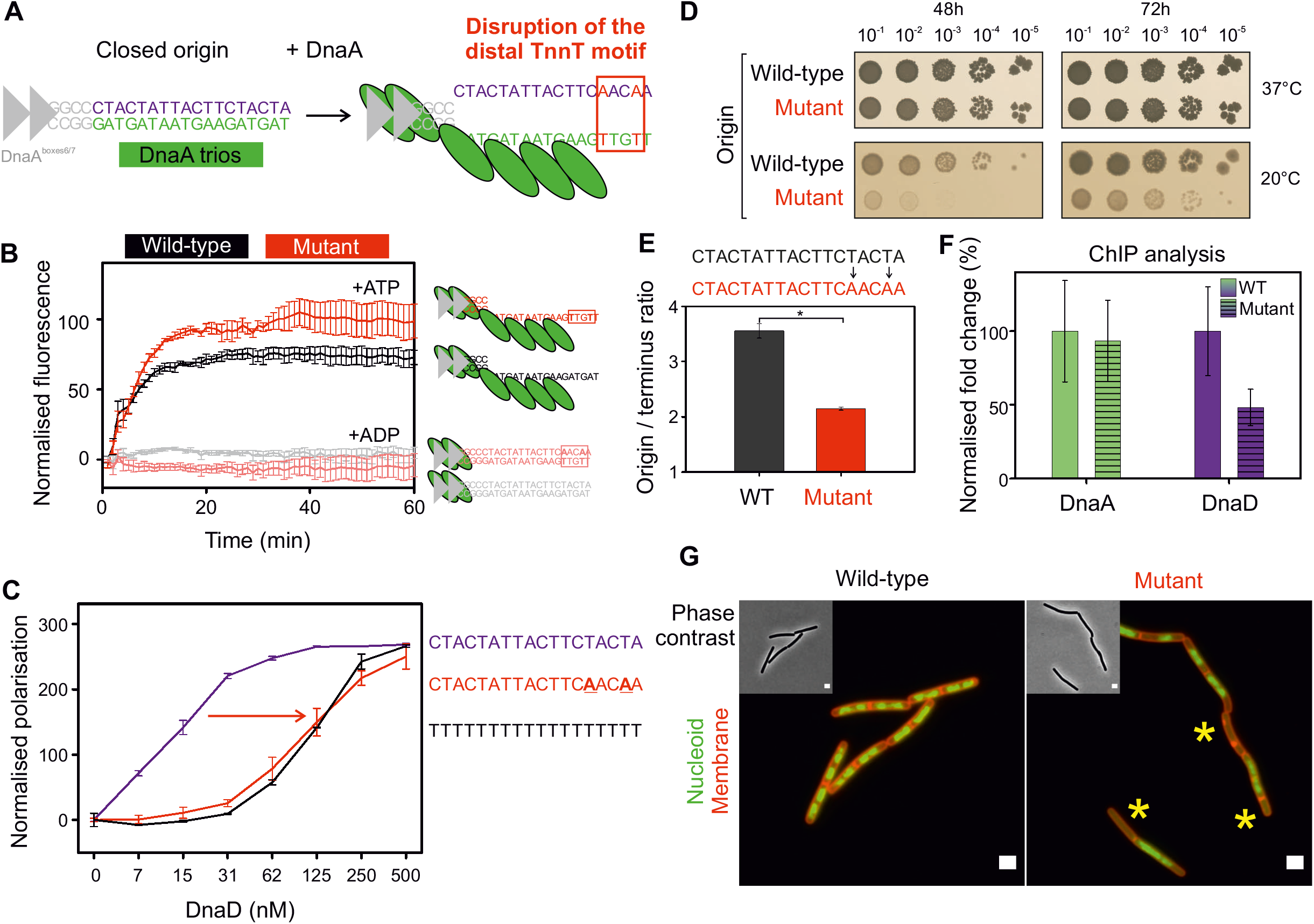
The DRE promotes specific ssDNA binding activity of DnaD *in vitro* and is required for efficient DNA replication initiation *in vivo*. **(A)** Schematics of *B. subtilis* origin unwinding region showing the potential impact of a mutation disrupting the distal 5’-TnnT-3’ element of the DRE with respect to DnaA boxes. **(B)** Strand separation assay showing that DnaA is comparably able to unwind substrates mimicking the wild-type origin or an origin where the distal DnaA trios have been disrupted as shown in panel (A). Background corresponding to the basal fluorescence of DNA complexes was subtracted from the curves. Grey and pink lines respectively show that DnaA does not unwind wild-type or mutant sequences in the presence of ADP. Black and red lines show that DnaA can unwind both wild-type and mutant origin complexes in the presence of ATP. Error bars indicate the standard error of the mean for 3-4 biological replicates. **(C)** Fluorescence polarisation analysis showing that a decrease in DnaD binding affinity to a DRE mutant where a 5’-TnnT-3’ motif has been mutated. The purple line (oCW1039) shows binding of DnaD to a sequence corresponding to the DRE. The red line (oCW1247) indicates that DnaD binding to a substrate where a 5’-TnnT-3’ element has been mutated is comparable to the profile of a same size polyT sequence (black line, oCW1088). **(D)** Spot titre analysis of the DRE mutant from panel (A) shows that introducing this mutation *in vivo* confers a cold-sensitive phenotype. Wild-type (168CA), Mutant (CW691). **(E)** Marker frequency analysis of the DRE mutant shown in panel (E) indicates that cells harbouring this mutation are impaired in DNA replication initiation at 20°C as measured by quantitative PCR. * shows a p-value of 0.018. Error bars show the standard error of the mean for 3 biological replicates. WT (168CA), Mutant (CW691). **(F)** ChIP analysis showing that the DRE mutant impairs the recruitment of DnaD to the origin. While DnaA is similarly recruited to the wild-type or mutant origins, DnaD was significantly less enriched in a background where a 5’-TnnT-3’ element of the DRE was mutated. The normalised fold change corresponds to the fold change in %IP observed for recruitment of DnaA and DnaD at the origin compared to background. Experiments were performed at 37°C and 100% indicates the reference fold change observed in wild-type cells. * shows a p-value of 0.0399 and n/s a non-significant difference. WT (168CA), Mutant (CW691). Primers used for the origin in panels (F-G) annealed within the *incC* region. Error bars show the propagated standard error of the mean for 3 biological replicates. **(G)** Representative microscopy images show that the mutation impairing the DRE leads to the appearance of anucleated cells at 20°C. Green fluorescence corresponds to imaging of the nucleoid by DAPI staining and red fluorescence depicts cellular membranes (Nyle red). Wild-type (168CA), Mutant (CW691). Scale bar indicates 2μm.

In contrast to the retained DnaA activities on substrates where the distal 5’-TnnT-3’ motif was changed to 5’-AnnA-3’, fluorescence polarisation *in vitro* revealed that these mutations significantly decreased DnaD binding (Fig. 6C). Together, these data suggest that disrupting the distal 5’-TnnT-3’ motif relative to the DnaA-boxes specifically impairs DnaD binding activity.

Next, we engineered a strain with the 5’-TnnT-3’ motif furthest from the DnaA-boxes mutated to 5’-AnnA-3’ (5’-CTACTATTACTTC<u>T</u>AC<u>T</u>A-3’ ➔ 5’-CTACTATTACTTC<u>A</u>AC<u>A</u>A-3’). It was observed that this mutant displays a growth defect at 20**°**C (Fig. 6D) and marker frequency analysis showed that the strain has a significantly lower DNA replication initiation frequency compared to wild-type cells at the lower temperature (Fig. 6E). Importantly, ChIP showed that DnaD recruitment to *oriC* was specifically reduced in the origin mutant, whereas the recruitment of DnaA was relatively unaffected (Figs. 6F and S10A-C). Consistent with a replication defect, fluorescence microscopy revealed that mutant cells had abnormal chromosome content, including an increase in cells lacking DNA at the restrictive temperature (Figs. 6G and S10D). Taken together, the results are consistent with the DRE functioning as a specific ssDNA binding site for DnaD within the *B. subtilis* chromosome origin unwinding region.

## DISCUSSION

Here we characterized the DNA binding activities of the essential *B. subtilis* DNA replication initiation protein DnaD. We found that DnaD binds a new ssDNA binding motif (DRE) within the *B. subtilis* chromosome origin via the DnaD^CTT^ (summarized in Figs. 7A-B). Based on the findings that DnaD tetramerization is required for maximal ssDNA binding (Fig. 3E-F), that the C-terminal tails likely extend away from the DnaD dimer:dimer interface (Fig 4C), and that the distance between a pair 5’-TnnT-3’ motifs is flexible (Fig. 5G), we speculate that at *oriC* each DnaD dimer (within a tetramer) donates one DnaD^CTT^ to engage the DRE (Fig. 7B). Moreover, based on preference for ssDNA (Fig. 5), we propose that DnaD engages the DRE following DNA strand separation by DnaA. These results further define the step-wise assembly pathway of DnaA and DnaD at the *B. subtilis* chromosome origin.

**Figure 7.**
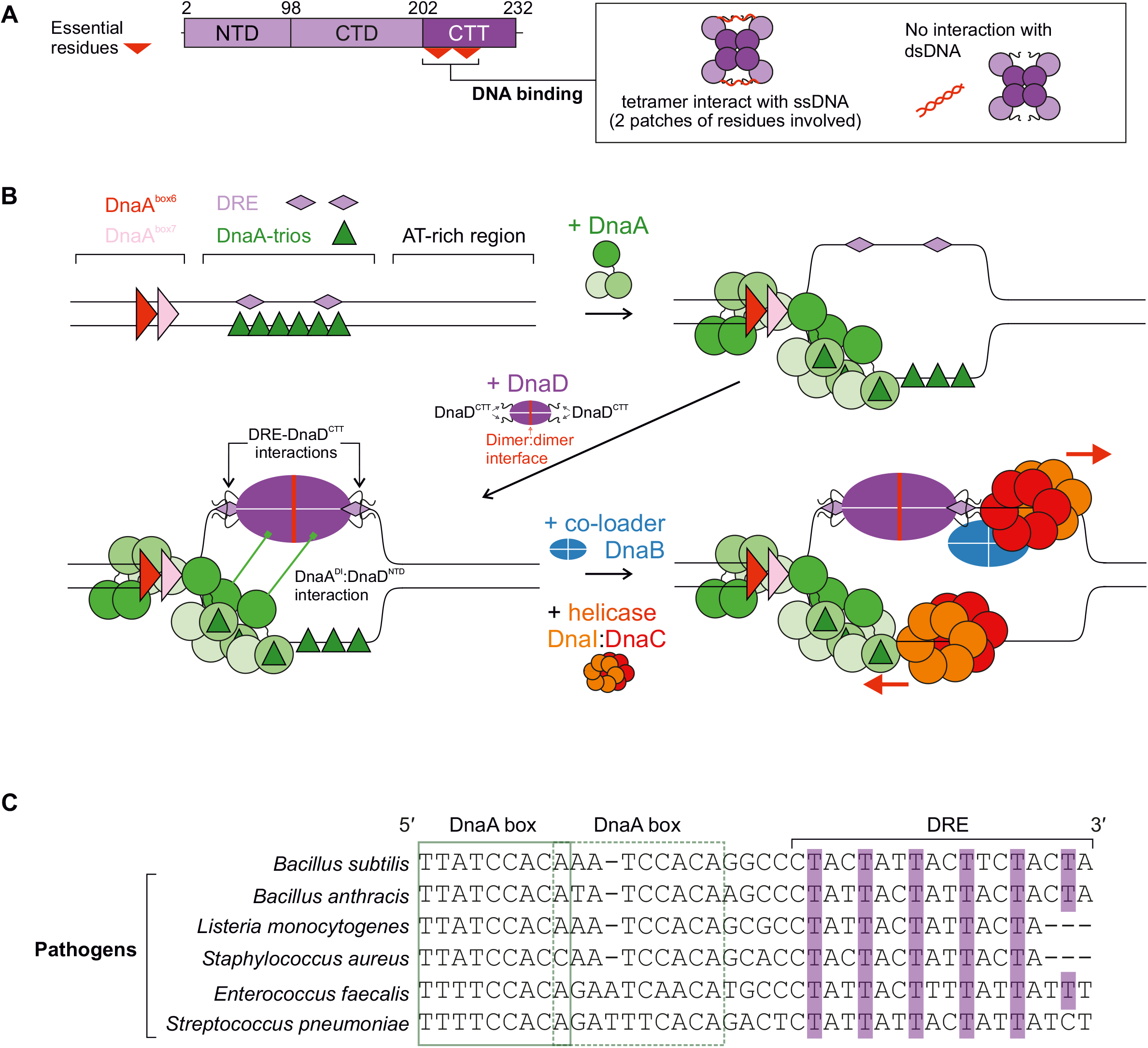
DnaD DNA binding activities at *oriC* culminate with it binding the DRE and leading to the onset of bidirectional replication. **(A)** DnaD mutations identified two patches of residues in the DnaD^CTT^ that are essential for ssDNA binding. **(B)** Model of the chromosomal replication initiation pathway in *B. subtilis*. DnaA binds DnaA-boxes in the *incC* region, leading to oligomer formation on DnaA-trios and initial opening of the origin. DnaD is then recruited and loaded on the exposed top strand (DRE) through both DRE:DnaD^CTT^ and interactions and DnaA domain I:DnaD N-terminal domain, which acts as a scaffold for further recruitment of the co-loader protein DnaB and bidirectional loading of the helicase complex DnaI:DnaC. DnaD C-terminal tail (DnaD^CTT^), DnaA domain I (DnaA^DI^), DnaD N-terminal domain (DnaD^NTD^). **(C)** Nucleotide alignment of DRE sequences showing the regular spacing and preservation of 5’-TnnT-3’ repeats (thymidines highlighted in purple) throughout the origins of *B. subtilis* and pathogens. The relative position of proximal DnaA boxes is indicated by green outlines.

### A mechanism for bidirectional DNA replication at a bacterial chromosome origin

A model was proposed for helicase loading at one end of a DnaA oligomer, where the AAA+ class of helicase loader engages the AAA+ motif of DnaA and guides deposition of helicase onto ssDNA (Fig. 7B) (18). This mechanism results in helicase loading around ssDNA in the correct orientation for 5′➔3′ translocation through *oriC*.

In contrast, the mechanism for loading a helicase onto the complementary strand was unclear. In *E. coli*, biochemical and genetic assays have suggested that DnaA domain I interacts directly with the helicase (59, 60). Moreover, it has been shown that DnaA^E21^ is essential for *E. coli* viability and required for helicase loading *in vitro* (61). In a recent study in *B. subtilis*, we identified residues in DnaA^DI^ that are essential for cell viability and required for the direct recruitment of DnaD to *oriC* (19). Generalizing, it appears that DnaA^DI^ acts as a critical protein interaction hub for helicase recruitment in diverse bacterial species, albeit through mechanisms involving interactions with distinct proteins. Importantly however, these protein:protein interactions alone do not resolve how a second helicase is recruited to a specific DNA strand with the correct geometry to support bidirectional replication.

Many bacterial chromosome origins encode a core set of sequence elements that direct DnaA oligomerization onto ssDNA, thus dictating the strand onto which the AAA+ helicase chaperone would dock (17). Here we report that the DRE, located opposite to where the DnaA oligomer binds, provides a mechanism for orchestrating strand-specific DnaD recruitment in *B. subtilis*; this mode of action is potentially conserved in many *Firmicutes* including the pathogens *Staphylococcus, Streptococcus, Enterococcus,* and *Listeria* (Fig. 7C) (62). Considered together with studies of primosome assembly at a single-strand origin *in vivo,* where binding of DnaD to ssDNA promotes subsequent helicase loading (63) (Fig. S11), we propose that the specific interaction of DnaD with the DRE provides a pathway for loading a second helicase to support bidirectional DNA replication (Fig. 8B).

### Outstanding questions for helicase loading mechanisms

The proposed model for DnaD recruitment raises several fundamental questions. How does DnaD (along with DnaA and DnaB) orientate the DnaI:helicase complex onto the strand encoding the DRE? We speculate that within the open complex formed at *oriC*, the distal junction between dsDNA and the unwound ssDNA could physically influence the subsequent events. Additionally, protein:protein interactions at *oriC* nucleoprotein complexes involving DnaA, DnaD and DnaB could play roles.

How is the temporal loading of two helicases orchestrated at *oriC*? Studies of *E. coli* helicase loading onto artificial DNA scaffolds that mimic an open chromosome origin indicated that DnaA preferentially recruits helicase onto the strand corresponding to where the DRE is located (64). Whether this order of recruitment holds during the physiological helicase loading reaction is unclear. While we favour a model where loading of the two helicases at *oriC* is reproducibly sequential, an alternative hypothesis is that loading of the two helicases is stochastic.

Are there other sequence elements within *oriC* that direct helicase loading? The discovery of the DnaA-trios and the DRE indicate that bacterial chromosome origins encode more information than previously appreciated. We note that many chromosome origins contain an intrinsically unstable AT-rich region (57, 65) where one of the helicases is loaded *in vitro* (66). The relative redundancy of dual 5’-TnnT-3’ repeats and the flexible distance separating these elements (Fig. 5G) supports the notion that additional sequence dependent information may be located within these AT-rich sites, or elsewhere. Further characterization of the nucleoprotein complexes formed at *oriC*, as well as dissection of downstream helicase loader proteins, will be needed to provide answers.

What about bacteria lacking a *dnaD* homolog (67, 68)? One hypothesis is that DnaA domain I itself could bind ssDNA and promote helicase loading. The structure of DnaA domain I is similar to that of the K homology (KH) domain (61), which is often observed as a module for binding single-stranded nucleic acids. In *E. coli* it has been reported DnaA domain I alone both specifically interacts with ssDNA from *oriC* and directly contacts the replicative helicase (59–61). We also speculate that there may be species specific ssDNA binding proteins that could act analogously to DnaD. Supporting this notion, the surface of DnaA domain I that interacts with DnaD appears to be the same surface used for non-homologous proteins to bind DnaA in other organisms (69–72).

## DATA AVAILABILITY

All plasmids and strains are available upon request. Microscopy data reported in this paper will be shared upon request. Raw 2D electron microscopy data is available from the Electron Microscopy Data (EMD-13663).

## ACKNOWLEDGEMENTS

SEC-MALS experiments were performed by Dr Andrew Leech at the Molecular Interaction laboratory as a service from the University of York.

## AUTHOR CONTRIBUTIONS

CW, SP, DS, SF, EM, NBC, PS, TRDC, AI, HM contributed to the conception/design of the work. CW, SP, NBC, AI generated results presented in the manuscript. NBC collected the cryo-EM data, TRDC and AI processed the cryo-EM data. CW, SP, AI created Figures. CW, HM, AI wrote the manuscript. CW, HM, SP, PS, AI edited the manuscript.

---

**Figure S1.**
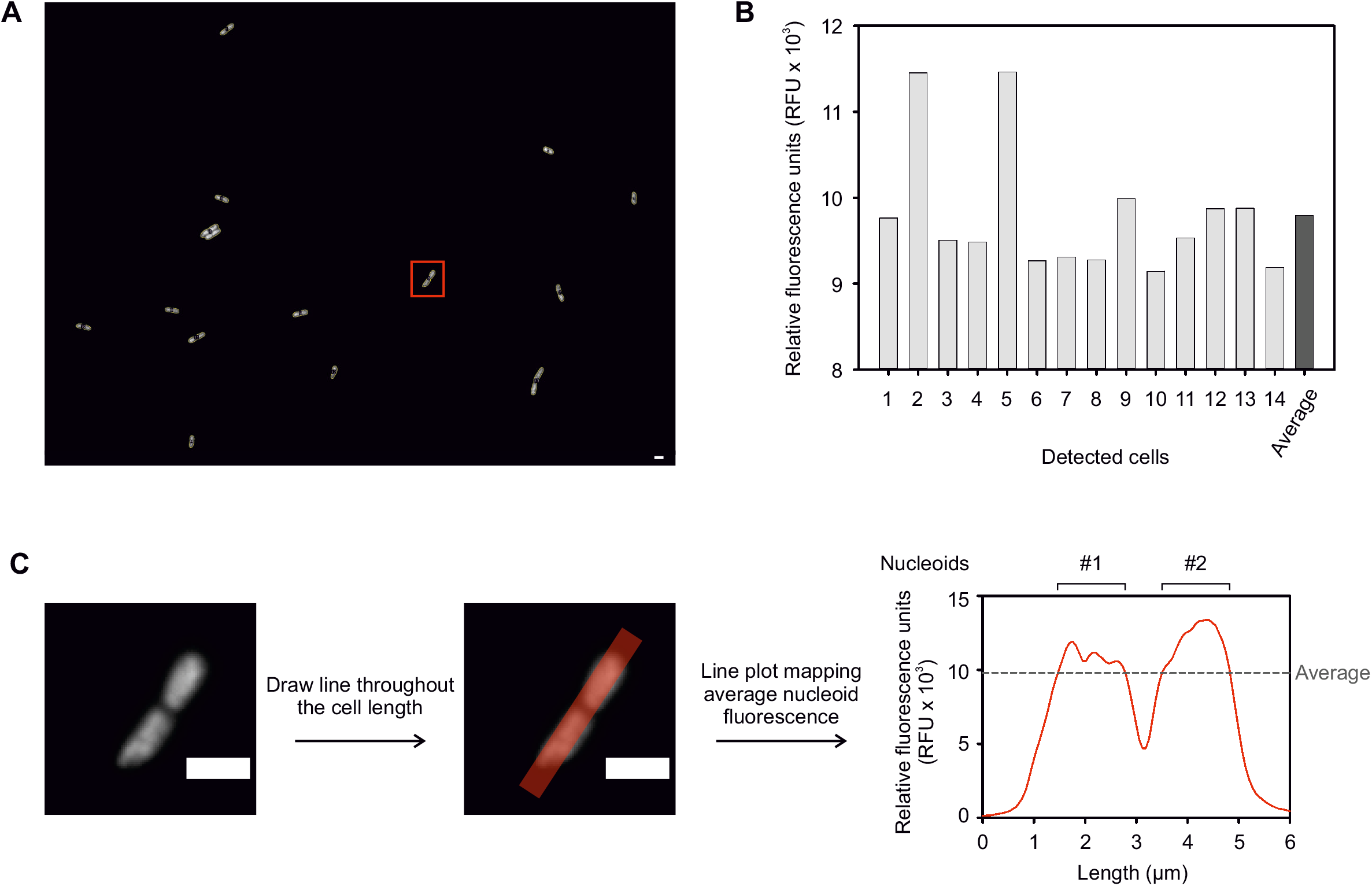
Microscopy analysis of the number of nucleoids per cell. **(A)** Detection of cells across an entire field of view using the ImageJ software. The red square indicates the cell studied in panel (C). **(B)** Mean of the fluorescence intensity for individual cells detected in panel (A) and average cytoplasmic fluorescence for this field of view. **(C)** Determination of the number of nucleoids at a single cell level. For each cell, a line was drawn across its longitudinal axis and the corresponding fluorescence intensity was plotted as a line graph. One nucleoid corresponds to a continuous area higher than the average cell fluorescence calculated in panel (B).

**Figure S2.**
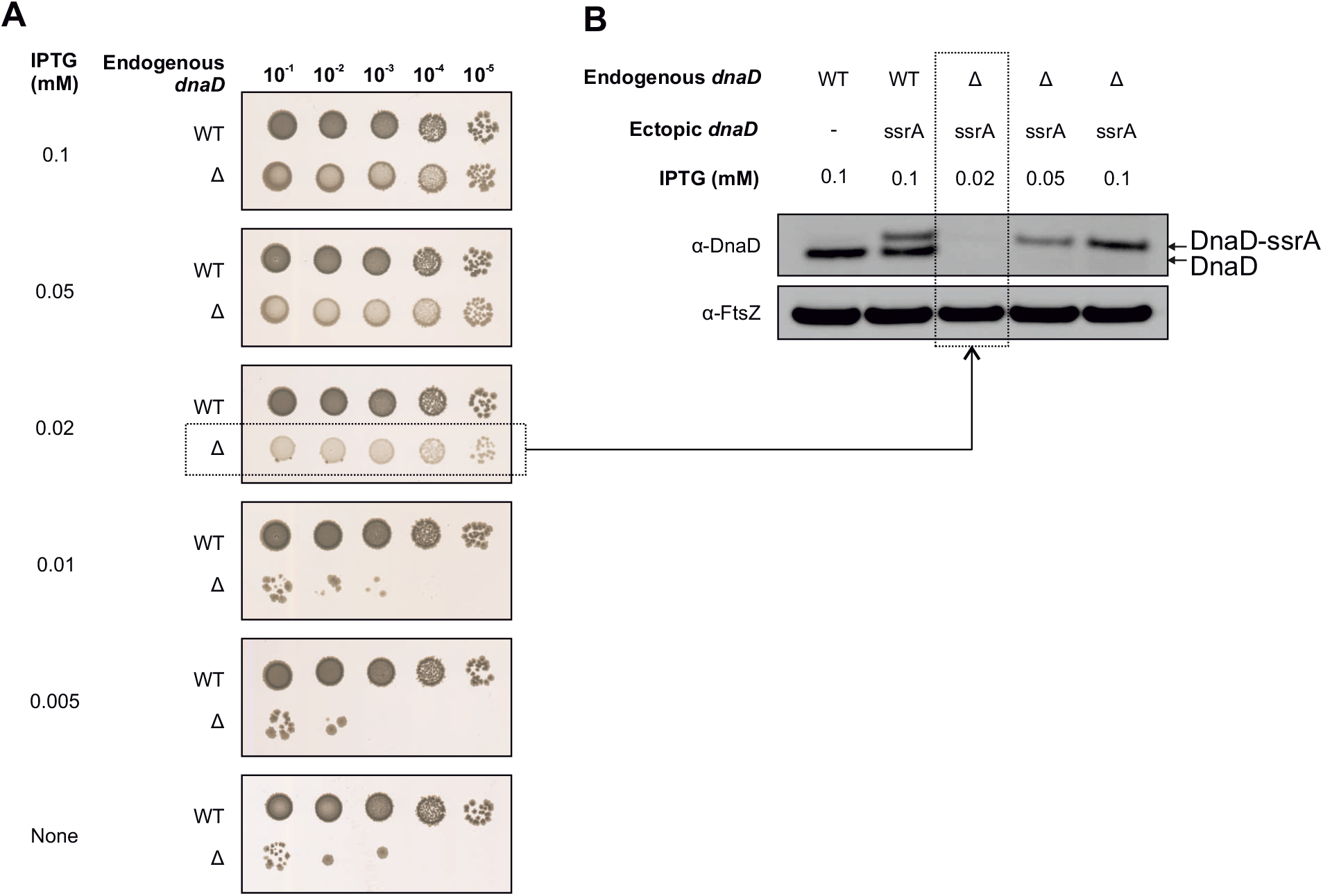
Low levels of DnaD expression sustain cell growth. **(A)** DnaD-SsrA was titrated via IPTG induction. The *dnaD-ssrA* cassette was able to sustain growth at IPTG concentration of 0.02 mM and above. WT (CW162), Δ (CW197). **(B)** Immunobloting shows that expression of DnaD was undetectable in viable colonies grown with 0.02 mM IPTG. The tubulin homolog FtsZ was used as a loading control.

**Figure S3.**
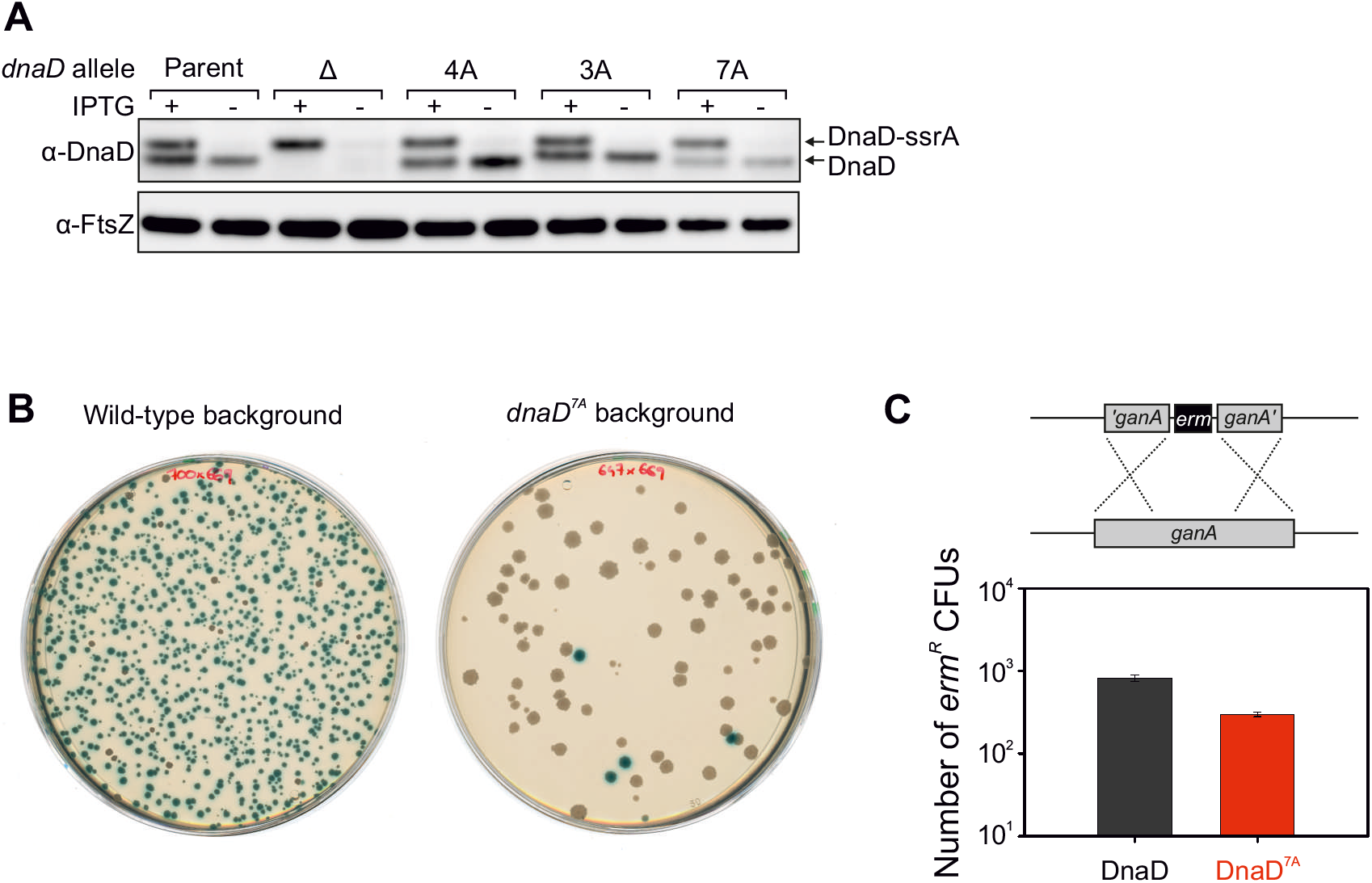
DnaD ssDNA binding characterisation. **(A)** Immunobloting of multiple alanine substitution DnaD variants targeting positively charged and aromatic residues within the DnaD^CTT^. The presence of IPTG shows expression of the ectopic DnaD-SsrA used for complementation. The tubulin homolog FtsZ was used as a loading control. Parent (Cw162), Δ (CW197), 4A (CW412), 3A (CW415), 7A (CW647). **(B)** Transformation plate showing a decrease in the number of blue colonies obtained in the *dnaD^7A^* background in an attempt to knock-out *dnaD-ssrA*. **(C)** Control DNA showing that strains transformed in panel (B) are both competent and integrated an *erm* cassette (*erm^R^* CFUs) at the ectopic locus *ganA* at a similar rate.

**Figure S4.**
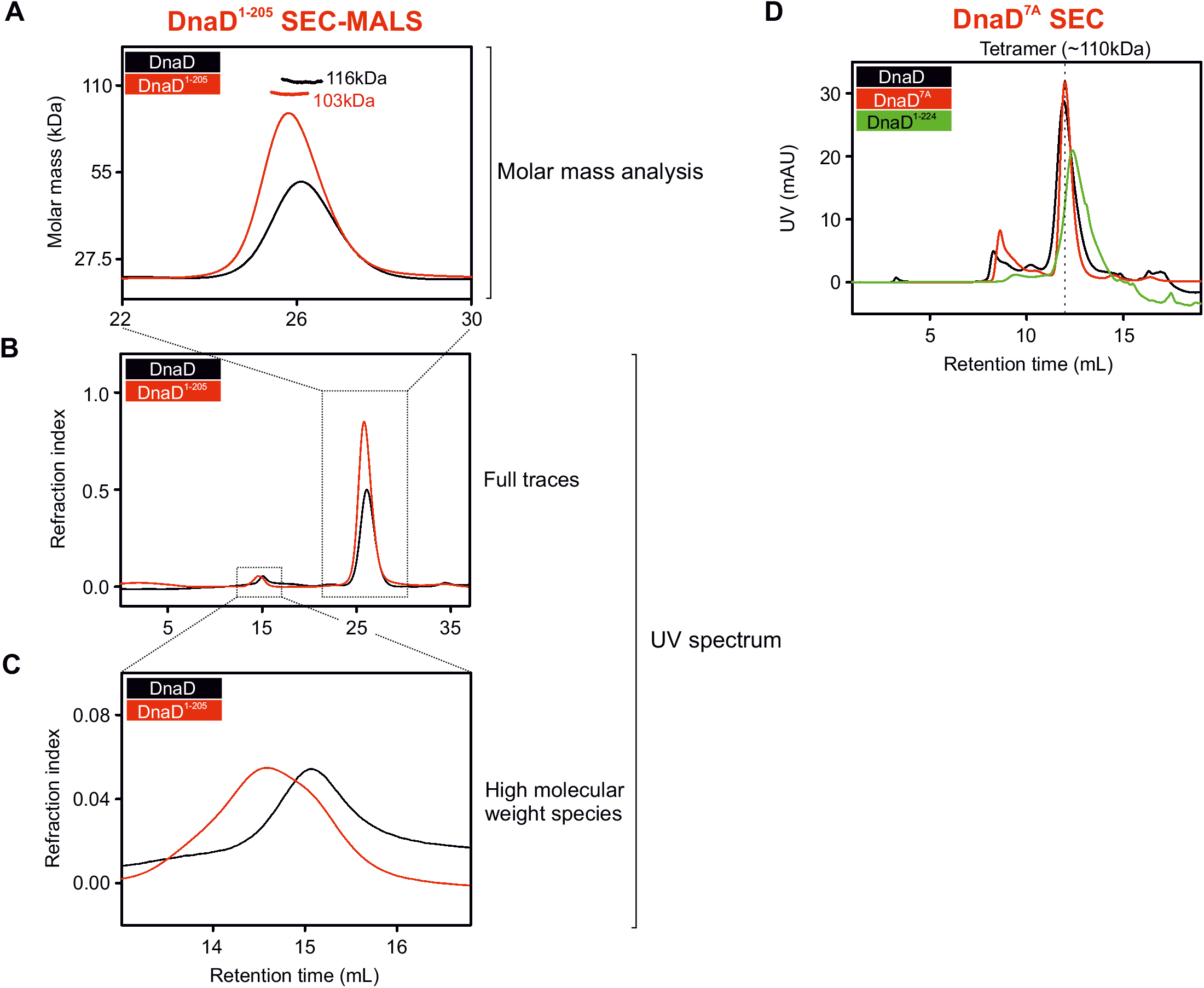
Size analyses of DnaD variants. **(A-C)** SEC-MALS analysis of purified DnaD protein variants. The wild-type protein (DnaD, black) or a truncation of the DnaD^CTT^ (DnaD^1-205^, red) were both resolved as tetrameric species. The UV spectrum (continuous lines) was normalised as a relative refraction index. **(A)** The molar mass corresponding to each protein is shown as shorter/thicker lines overlapping the different peaks of the refraction index. Masses corresponding to each peak are annotated on the plot. **(B-C)** Full trace of the UV-spectrum represented as a refraction index during the SEC experiments showing that the DnaD variants did not display major fractions of aggregates (high molecular weight species). **(D)** SEC analyses of DnaD wild-type and DnaD^7A^ and DnaD^1-224^ show that the 7A allele and C-terminal truncation had a similar UV-spectrum to the wild-type protein. Both proteins were able to form tetrameric species (dotted vertical line).

**Figure S5.**
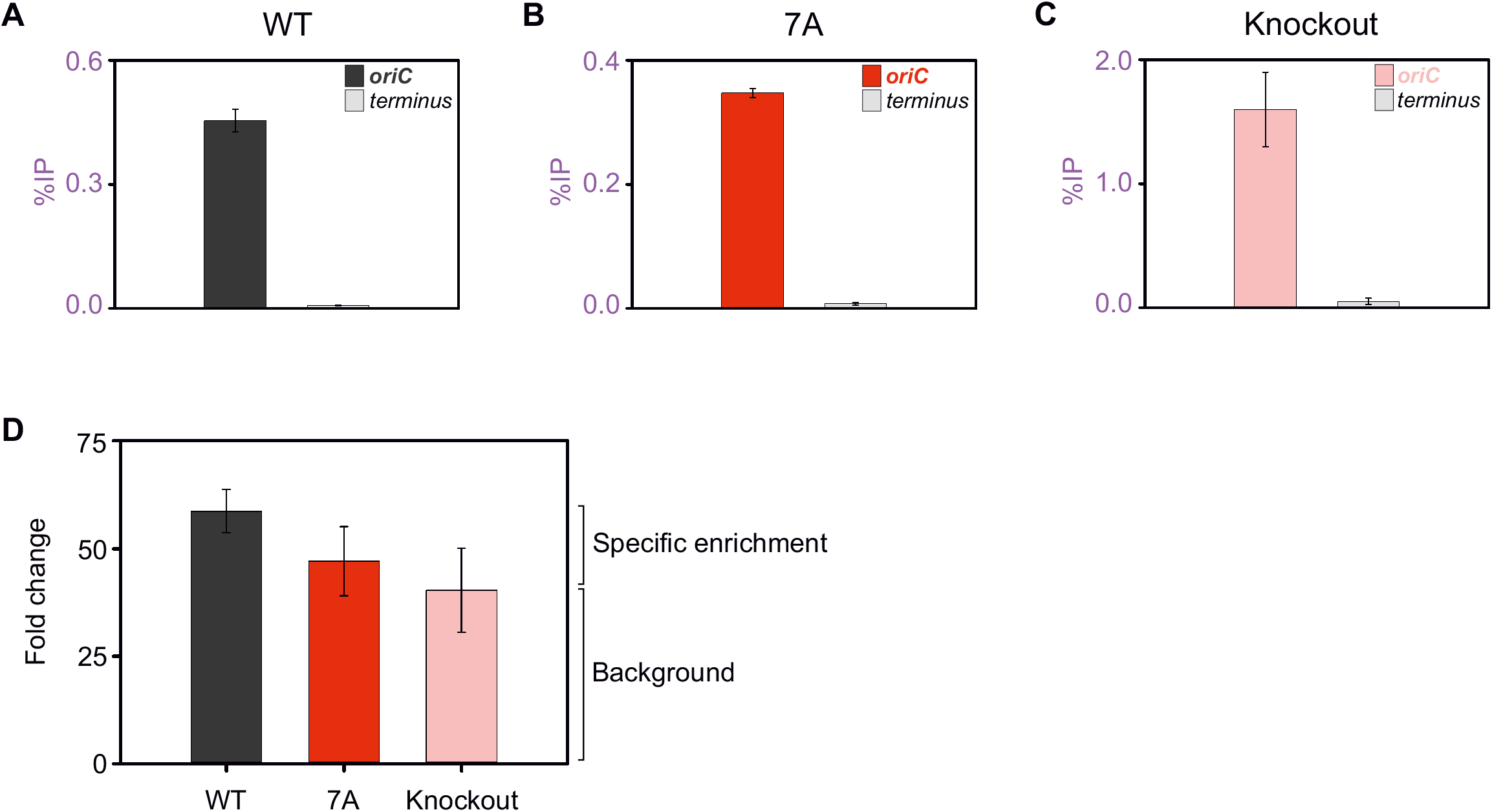
ChIP analysis of DnaD variants. **(A-C)** show the %IP detected for the recruitment of DnaD variants to the origin of replication for individual strains that also harboured the *dnaD-ssrA* cassette. **(A)** shows recruitment of DnaD in a wild-type background (Cw162). **(B)** shows recruitment of DnaD in the *dnaD^7A^* background (CW647). **(C)** shows recruitment of DnaD in a knockout strain of the endogenous *dnaD* (CW197). Primers used to amplify the origin in panels (A-C) annealed within the *incC* region. Error bars in panels (A-C) show the standard error of the mean for three biological replicates. **(D)** shows the fold change corresponding to the DnaD %IP detected in panels (A-C). The knockout condition shows that DnaD-SsrA (the only copy of DnaD in this strain) accounts for a significant amount of the IP detected in these experiments. Strains are the same as in panels (A-C) and error bars indicate the propagated error of the mean for averages shown in panels (A-C).

**Figure S6.**
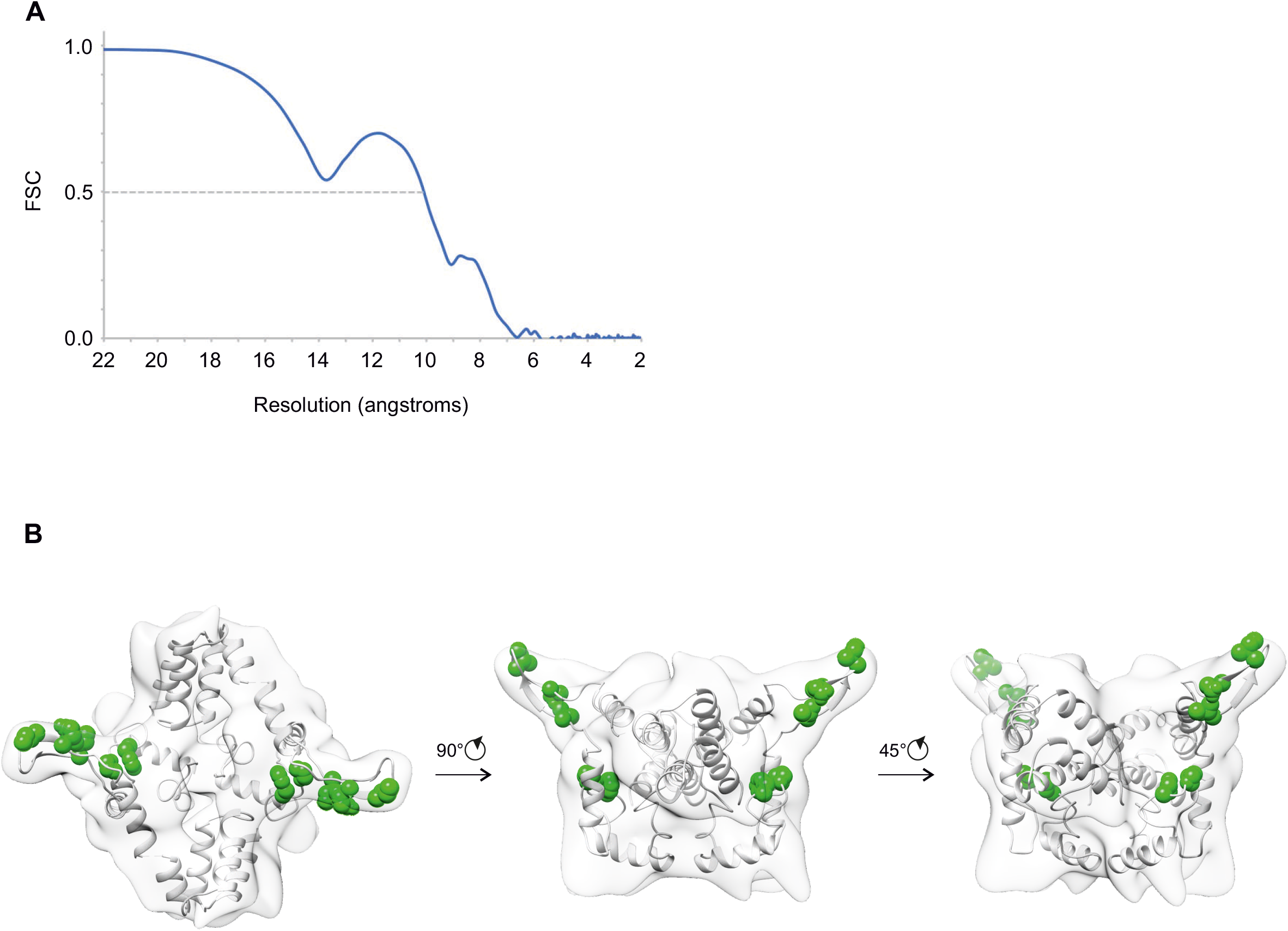
DnaD structure solved by cryo-EM. **(A)** Overall resolution of the DnaD dimer, derived from two independently refined half-maps, using the FSC=0.5 criteria. **(B)** Cryo-EM density map fitted with the available DnaD structures (N-terminal domains from PDB 2V79 and C-terminal domains from 2ZC2) with amino acid residues involved in binding DnaA shown in green.

**Figure S7.**
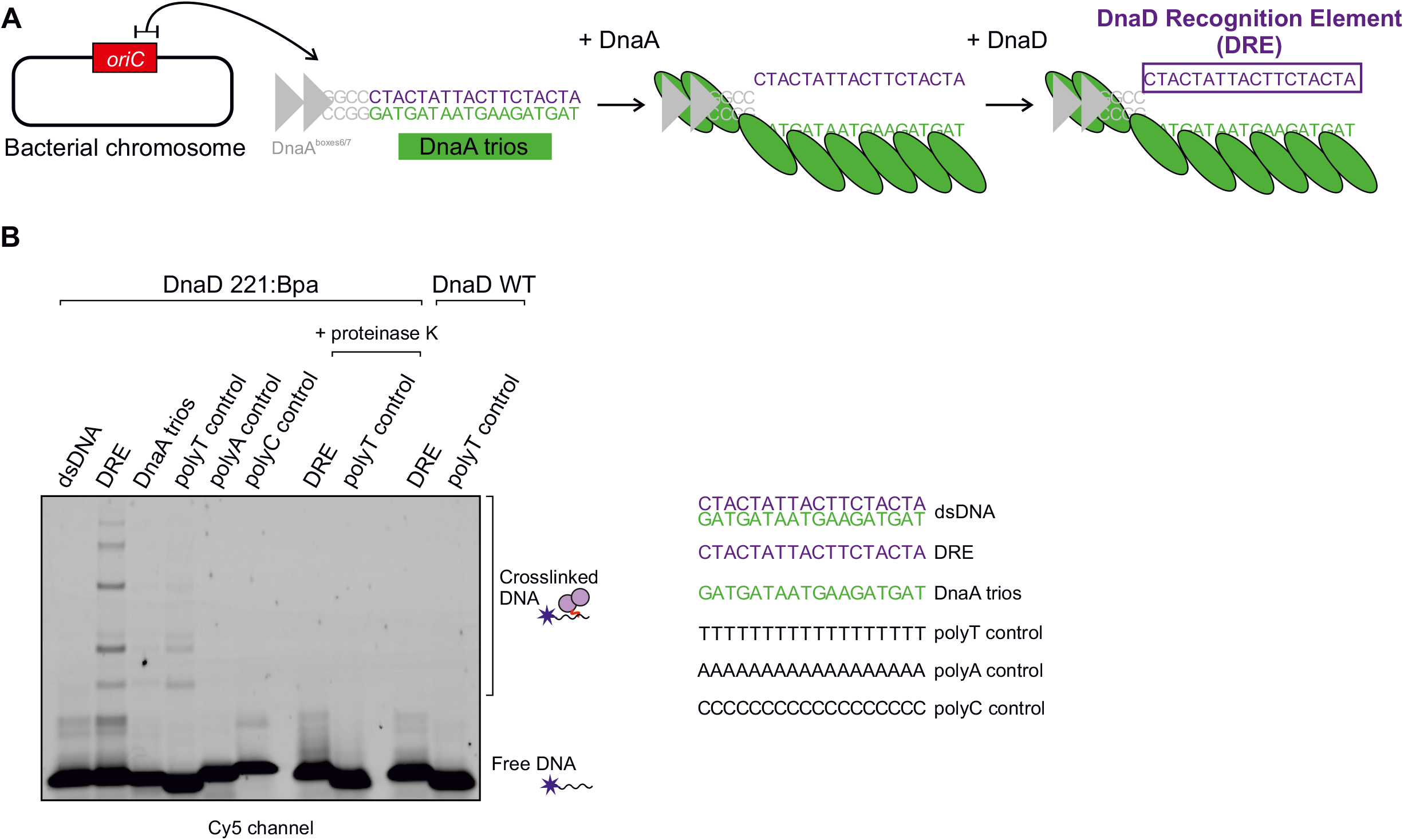
Crosslinking of DnaD to ssDNA substrates. **(A)** Illustration of the proposed basal origin unwinding mechanism in *B. subtilis*. The closed origin region *incC* is bound by DnaA on DnaA boxes, which leads to partial melting of the origin and strand separation via DnaA oligomer formation on the DnaA-trios. The complementary sequence to the DnaA-trios is proposed to be a specific binding substrate for DnaD. **(B)** Bpa crosslinking assay showing that DnaD interacts with the DRE via the amino acid residue 221 in the DnaD^CTT^. Incubation with Cy-5 labelled oligonucleotides shows that DnaD 221:Bpa does not crosslink with a dsDNA complex formed by the DRE and DnaA-trios. When used as ssDNA substrates, DnaD 221:Bpa specifically crosslinked the DRE. Treatment via proteinase K shows that binding of DnaD 221:Bpa to ssDNA was specific. Incubation with wild-type DnaD shows that the native protein was not able to be crosslinked onto ssDNA. Oligonucleotide sequences are indicated to the right of the gel: dsDNA (oSP1132:oCW1040), DRE (oSP1132), DnaA trios (oSP1133), polyT control (oSP1135), polyA control (oSP1137), polyC control (oSP1134).

**Figure S8.**
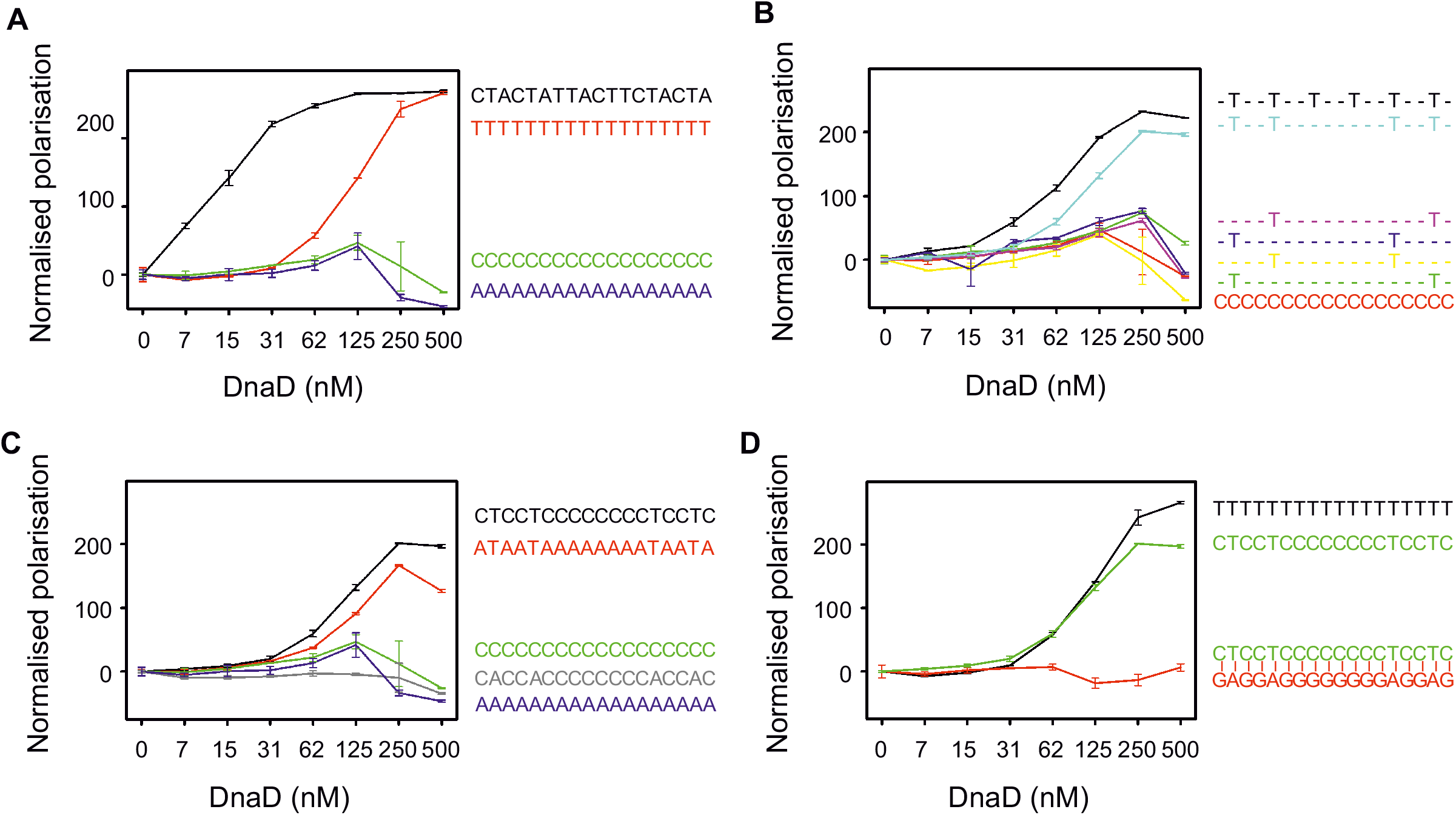
Two 5’-TnnT-3’ repeats are required for DnaD ssDNA binding *in vitro*. **(A-D)** show fluorescence polarisation analyses of DnaD binding to a range of DNA substrates. Error bars in show the standard error of the mean for 2-5 biological replicates. **(A)** shows binding to homopolymers of size and sequence comparable to the DRE. The black line shows binding to a sequence corresponding to the DRE (oCW1039), the red line to a polyT18 ssDNA (oCW1088), the green line to a polyC18 substrate (oCW1089) and the blue line to a polyA18 (oCW1090). **(B)** shows DnaD binding to 5’-TnnT-3’ motifs located within an inert ssDNA substrate and that all thymidines equally contribute to ssDNA binding. Black (oCW1139), cyan (oCW1128), pink (oCW1167), blue (oCW1166), yellow (oCW1155), green (oCW1127) and red (oCW1089). **(C)** shows that DnaD binding is specific to 5’-TnnT-3’ motifs located within otherwise inert ssDNA substrates, and that dual repeats of 5’-AnnA-3’ motifs are not recognised. Black (oCW1128), red (oCW1129), green (oCW1089), grey (oCW1216) and blue (oCW1090). **(D)** shows that DnaD only binds a 5’-TnnT-3’ dual repeat in the context of ssDNA. Black polyT18 control (oCW1088), green 5’-TnnT-3’ dual repeat as ssDNA (oCW1128), red 5’-TnnT-3’ dual repeat as dsDNA (oCW1128:oCW1527).

**Figure S9.**
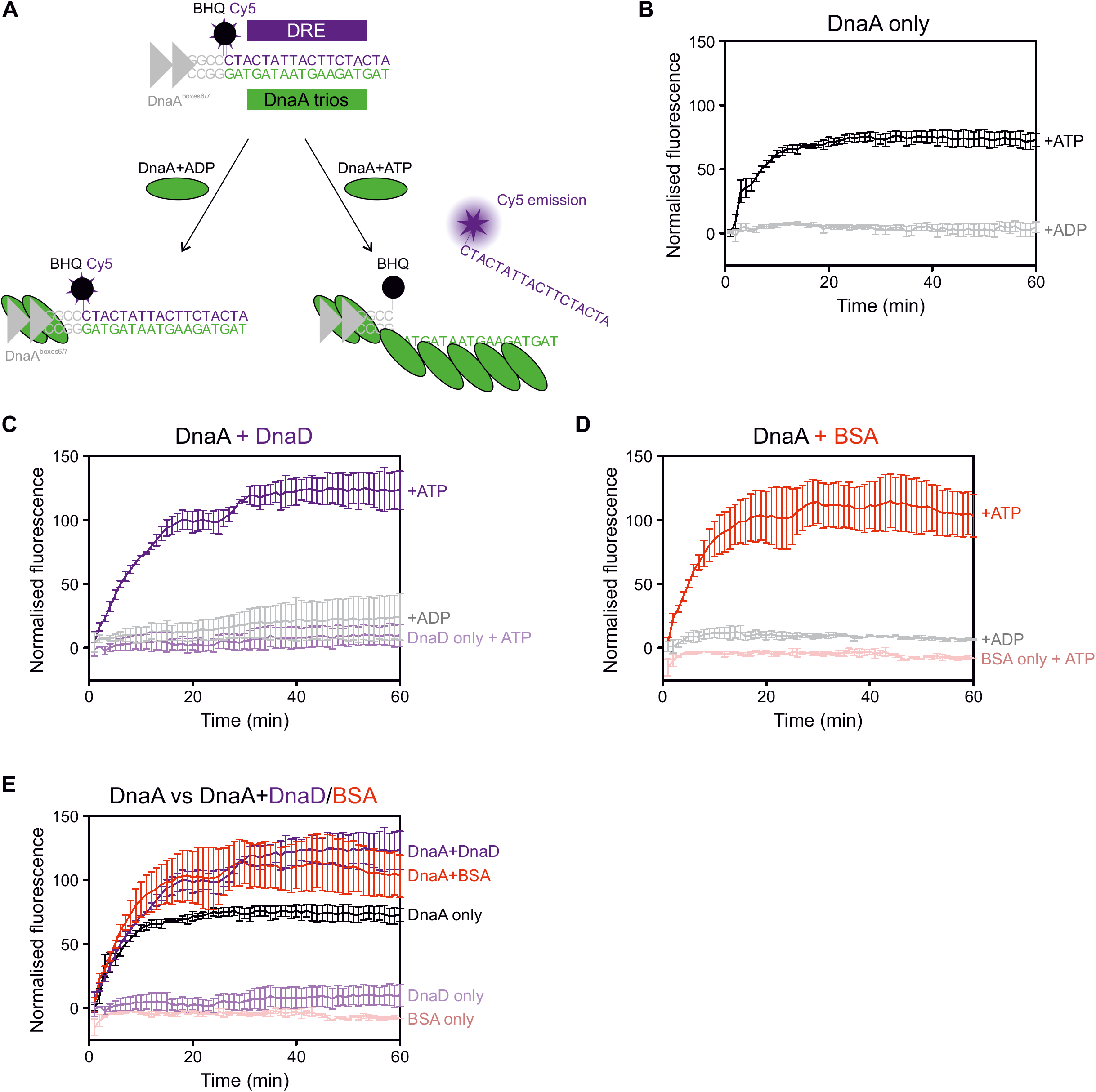
DnaD does not specifically contribute to substrate unwinding via DnaA. **(A)** Illustration of the strand separation assay setup used to detect DnaA-directed unwinding of DNA substrates. Three oligonucleotides are annealed to mimick the *B. subtilis* origin unwinding region including DnaA boxes (dsDNA binding via DnaA) and the DnaA trios/DRE region. The bottom strand is continuous and unlabelled. The top strand corresponding to the DnaA boxes and GC-rich region is labelled with a black-hole quencher (BHQ) at the 3’-end and the DRE is labelled with Cy5 at the 5’-end. As a fully dsDNA probe, the BHQ quenches fluorescence emitted by the Cy5 group. Upon incubation with DnaA and ADP, DnaA binds DnaA boxes, cannot engage the DnaA trios and no fluorescence remains quenched. In the presence of ATP, DnaA binds DnaA boxes and forms an oligomer on the DnaA trios, thereby displacing the DRE probe and allowing emission of Cy5 fluorescence. **(B-E)** Strand separation assays performed with the same probe in the presence of various proteins. DNA substrate: oHM558/oHM778:oHM590. Background corresponding to the basal fluorescence of the DNA probe was subtracted from the curves. Error bars show the standard error of the mean for three biological replicates. **(B)** shows that DnaA only separates strands in the presence of ATP. **(C)** shows that DnaD does not separate strands on its own in the presence of ATP, and that DnaA can still unwind substrates when incubated with DnaD and ATP. **(D)** shows that BSA (control protein) does not separate strands on its own in the presence of ATP, and that DnaA can still unwind substrates when incubated with BSA and ATP. **(E)** shows that strand separation is comparable when using DnaD or BSA along DnaA in the presence of ATP. DnaD or BSA cannot separate strands without DnaA.

**Figure S10.**
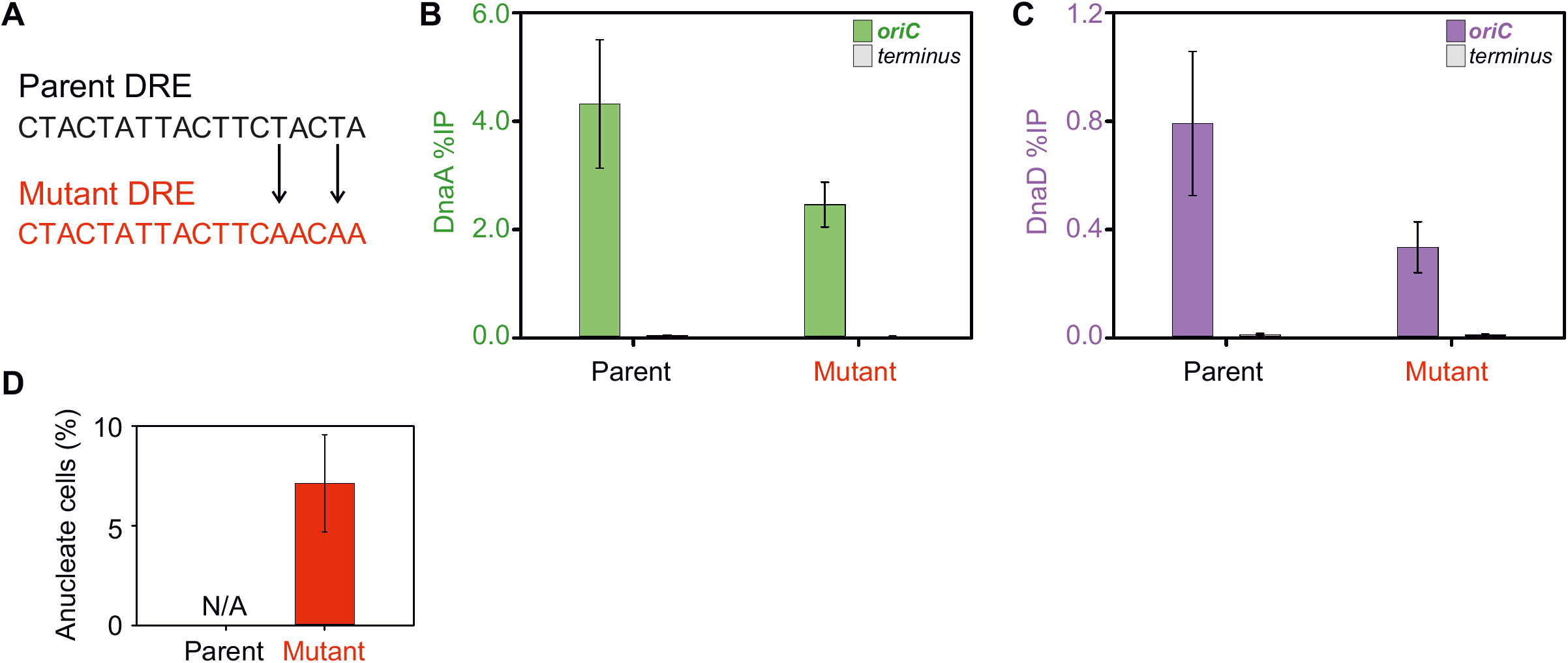
*In vivo* analysis of origin variants. **(A)** shows the mutation that was introduced in the DRE *in vivo.* The last 5’-TnnT-3’ motif present in the parent sequence was mutated to 5’-AnnA-3’. **(B-C)** show the %IP detected for the recruitment of DnaA and DnaD to the origin of replication variants using ChIP. Parent indicates a wild-type strain (168CA) and mutant a strain harbouring the mutation depicted in panel (A) (CW691). **(B)** shows recruitment of DnaA and **(C)** shows recruitment of DnaD. Primers used to amplify the origin in panels (B-C) annealed within the *incC* region. Error bars in panels (B-C) show the standard error of the mean for three biological replicates. **(D)** shows that the origin mutant displayed in panel (A) produces anucleate cells. Quantification was performed from two biological repeats, where over 750 cells were counted for each strain. N/A indicates that no anucleate cells were found and the error bar indicates the standard error of the mean.

**Figure S11.**
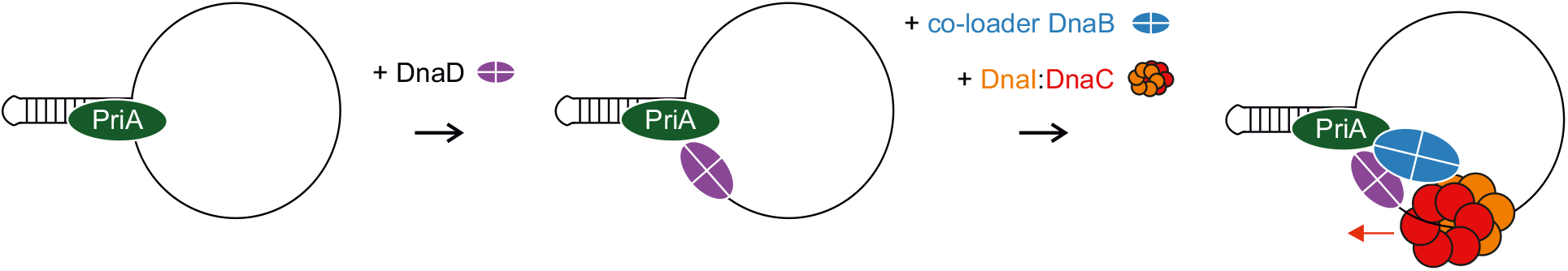
Model for helicase recruitment and loading in *B. subtilis* during PriA-dependent replication restart at a single-strand origin (*sso)*.

